# Changes in Active Site Loop Conformation Relate to the Transition toward a Novel Enzymatic Activity

**DOI:** 10.1101/2023.05.22.541809

**Authors:** Pauline Jacquet, Raphaël Billot, Amir Shimon, Nathan Hoekstra, Céline Bergonzi, Anthony Jenks, Eric Chabrière, David Daudé, Mikael H. Elias

**Author notes:** Correspondence: For the biochemical studies; For the structural studies. Current address: Univ. Grenoble Alpes, CNRS, CEA, IBS, F-38000 Grenoble.

## Abstract

Enzymatic promiscuity, the ability of enzymes to catalyze multiple, distinct chemical reactions, has been well documented and is hypothesized to be a major driver for the emergence of new enzymatic functions. Yet, the molecular mechanisms involved in the transition from one activity to another remain debated and elusive. Here, we evaluated the redesign of the active site binding cleft of the lactonase *Sso*Pox using structure-based design and combinatorial libraries. We created variants with largely improved catalytic abilities against phosphotriesters, the best ones being > 1,000-fold better compared to the wild-type enzyme. The observed shifts in activity specificity are large, ∼1,000,000-fold and beyond, since some variants completely lost their initial activity. The selected combinations of mutations have considerably reshaped the active site cavity *via* side chain changes but mostly through large rearrangements of the active site loops, as revealed by a suite of crystal structures. This suggests that specific active site loop configuration is critical to the lactonase activity. Interestingly, analysis of high-resolution structures hints at the potential role of conformational sampling and its directionality in defining an enzyme activity profile.

## 1. INTRODUCTION

Over the past years, it has become evident that enzymes may accept more than one substrate or catalyze more than one specific reaction, and much attention has been paid to investigating the role of enzyme promiscuity in the emergence of new catalytic functions.^1–3^ Promiscuous activities may confer adaptability and could constitute an advantage under selection pressure.^4, 5^ The transition between phenotypes has been shown to be relatively close in the genotype space, the acquisition of new functions being gradual and smooth in the course of evolution.^6, 7^ Catalytic promiscuity may therefore be used as a starting point for directed evolution strategies.^8–10^

The hypothesized recent emergence of phosphotriesterase (PTE) enzymes is a striking example of the role of enzyme promiscuity on selective adaptation. Indeed, the wide use of organophosphorus insecticides (OP) since the 1950s for agricultural purposes may have led to the emergence of highly efficient OP-degrading enzymes for the mobilization of the poorly-available phosphate.^11^ Several highly proficient bacterial PTE appear to have emerged from a promiscuous lactonase template belonging to the Phosphotriesterase-like Lactonase (PLL) family.^11, 12^ PLLs are enzymes naturally able to catalyze the hydrolysis of lactones, especially acyl-homoserine lactones involved in bacterial communication referred to as quorum sensing.^11, 13^ Beside their lactonase activity, most PLL show a promiscuous activity towards phosphotriesters, that may rely on the similarity of the tetrahedral intermediate involved in their respective reactions. PLL and PTE share a (β/α)_8_ fold with highly mobile loops that contribute to broaden their promiscuity to other substrates such as aryl-esters.^14^ Bacterial PTEs, although naturally efficient for the degradation of OP insecticides, have been further engineered *in vitro* for the development of biocatalysts close to catalytic perfection for decontaminating insecticides or chemical warfare agents.^15–22^ These variants may find application in bioremediation of agricultural contaminations or for prophylaxis protection against OP poisoning.^23, 24^ However, most PTE have been isolated from mesophilic microorganisms and show moderate stability limiting their biotechnological and medical uses.^25^ A subfamily of PLL, namely PLL-A, is composed of enzymes isolated from thermostable and hyperthermostable bacteria or archaea. Special attention has thus been paid to PLL-A for developing high-potential biocatalysts as robust enzymes that may offer numerous biotechnological advantages including resistance to high temperatures, tolerance to solvents, denaturants and long-term storage.^26^ Moreover, thermostable enzymes are relevant for directed evolution strategies as their robustness may buffer deleterious mutations and offer new evolutionary trajectories.^27–30^ In that context, the enzyme *Sso*Pox isolated from hyperthermophilic archaeon *Saccharolobus solfataricus* was investigated. *Sso*Pox is a natural lactonase with a promiscuous paraoxonase activity that has been considered due to its tremendous thermostability (T_m_ = 106 °C).^31–33^ This enzyme was previously engineered by our group and others to alter its enzymatic profile, including increase its phosphotriesterase activity. Such improved variants were previously shown to decontaminate a variety of phosphotriesters, such as insecticides, chemical warfare agents (CWA), or their surrogates and could decrease OP-poisoning in animal models.^34–40^

Here, we report an attempt to redesign *Sso*Pox’s active site using structure-based design and combinatorial strategies to alter its activity and understand the determinants for its activity profile (**Figure 1**). A structure-based design approach included comparisons between the structures of *Sso*Pox and the PTE from *Brevundimonas diminuta* to identify the key positions involved in the binding of substrates. A combinatorial library was synthesized using degenerated oligonucleotides to diversify these positions and an activity-based screening procedure was used to purge deleterious mutations. Mutants were then further screened against various substrates including lactones, esters, and phosphotriesters to identify improved variants and provide deep insights of their substrate promiscuity. The most efficient variants were produced and purified, and their catalytic parameters and melting temperatures were determined. The best variants show improvement of up to 1,000-fold in phosphotriesterase activity as compared to the wild-type enzyme. Remarkably, some of these variants show a specificity shift, namely simultaneous increase of phosphotriesterase and a decrease of lactonase activities of a combined ∼1,000,000-fold. This increased phospotriesterase activity translates into an improved ability to protect recombinant human acetylcholinesterase (rHAChE) from inhibition by OP. The determination of the tridimensional structures for some of the most improved variants reveals potential structural determinants for the observed transition in catalytic activity.

**Figure 1.**
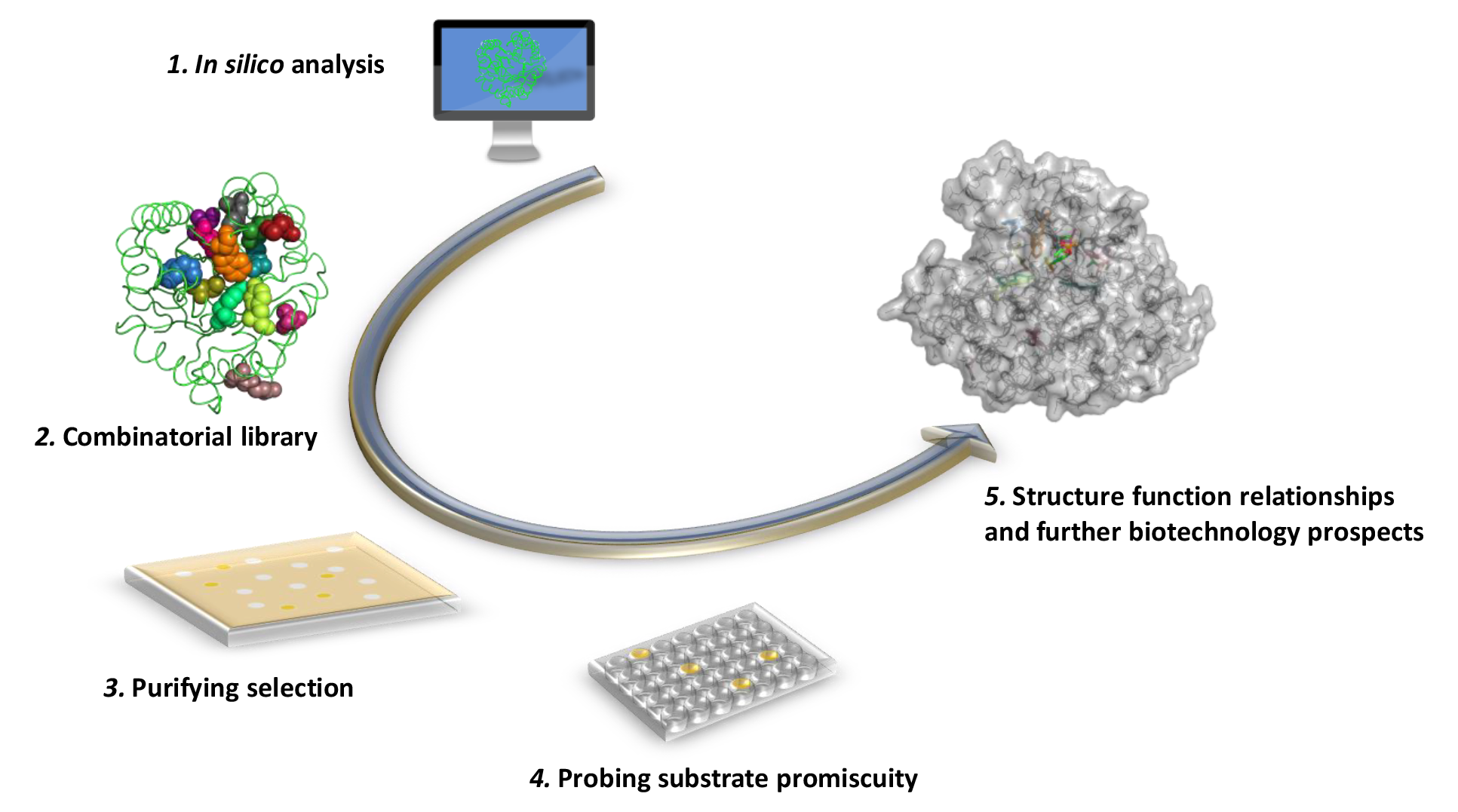
Structure-based design and directed evolution of *Sso*Pox. (1) *In silico* analyses were performed to compare *Sso*Pox with bacterial PTE from *Brevundimonas diminuta*. (2) 12 residues were targeted for designing a combinatorial library using a degenerated oligonucleotide-based strategy. (3) A solid-based screening was adapted to identify variants retaining a paraoxonase activity. (4) A miniaturized procedure was used to screen a 430-variant library against 9 substrates. (5) The most relevant variants were produced purified and characterized for further characterization.

## 2. MATERIAL AND METHODS

### Chemicals

Organophosphorus compound analogues were synthesized by Enamine Ltd. All other chemicals used for kinetic assays were ordered from Sigma-Aldrich.

### Design and Screening of the Combinatorial Library

Three dimensional structures of *Sso*Pox (PDB ID 2VC5) and *Bd*PTE (PDB ID 1DPM) were superimposed with PyMOL^41^ based on their (α/β)_8_ topology similarity. Thirteen residues were targeted for the construction of the combinatorial library, V27, L72, Y97, Y99, R154, T177, R223, L226, L228, C258, W263, W278 and I280. A randomized strategy based on degenerated oligonucleotides was applied for creating genetic diversity. The plasmid library was synthesized by GenScript, using degenerated oligonucleotides procedure to create a randomized and degenerated library. The desired mutations and the corresponding codon are described in **Table S1**. Then *Sso*Pox gene was cloned in pET32b-Δtrx vector using *NdeI* and *XhoI* restriction sites.

### Screening of the Library

The plasmid library was transformed into *Escherichia coli* strain BL21(DE_3_)-pGro7/GroEL (TaKaRa). Cells were grown overnight at 37°C on nitrocellulose membranes (Amersham™ Protran™, GE Healthcare) in plates containing ZYP agar medium (supplemented with 100 µg.mL^-1^ ampicillin and 34 µg.mL^-1^ chloramphenicol). For the induction, nitrocellulose membranes were transferred to ZYP agar medium plates (supplemented with 100 µg.mL^-1^ ampicillin and 34 µg.mL^-1^ chloramphenicol) containing L-arabinose 0.2% and CoCl_2_ 0.2 mM. Plates were incubated at room temperature overnight. Cell lysis was realized by thermal shock, with three cycles of 30 seconds at 37°C and -20°C. For the selection of clones, nitrocellulose membranes were transferred on plates containing ethyl-paraoxon 250 µM, agarose 1.5%, and *activity buffer* (NaCl 150 mM, HEPES 50 mM, CoCl_2_ 0.2 mM pH 8) **(Figure S1)**. Colonies capable of degrading paraoxon were identified by their yellow color through visual inspection of the plate, and were isolated. A total of 430 variants was obtained.

The selected variants were then expressed in liquid medium as previously described.^36^ Shortly, variants were grown on 96-well plates in LB medium (supplemented with 100 µg.mL^-1^ ampicillin and 34 µg.mL^-1^ chloramphenicol) overnight at 37°C. 96-well plates containing 1 mL of ZYP medium were inoculated. After 5 hours of growth at 37°C, 0.2% L-arabinose and CoCl_2_ 0.2 mM were added for induction and cultures were grown overnight at room temperature. Cells were harvested by centrifugation and lysis was performed in *lysis buffer* (50 mM HEPES pH 8.0, 150 mM NaCl, 0.2 mM CoCl_2_, 0.25 mg.mL^-1^ lysozyme, 0.1 mM PMSF and 10 µg.mL^-1^ DNAseI). Lysates were partially purified by heating at 70°C for 20 min.

Activity assays were performed on partially purified variants for nine substrates: ethyl-paraoxon and p-nitrophenol acetate (250 µM, 405 nm), for the tested phosphotriester analogues (10 µM, fluorescence 360/40, 460/40 nm), and for 3-oxo-C10 and 3-oxo-C12 HSL (250 µM, 577 nm) **(Figure S2)**. Twelve variants were selected, sequenced by GATC Biotech™ and purified.

### Production and Purification of *Sso*Pox Wild-Type and Variants

*Sso*Pox variants were produced in *E. coli* BL21(DE_3_)-pGro7/GroEL (TaKaRa) chaperone expressing strain using a pET32b-Δtrx plasmid.^31^ Cultures were performed in auto-inducible ZYP medium (supplemented with 100 µg.mL^-1^ ampicillin and 34 µg.mL^-1^ chloramphenicol), induction took place when OD_600 nm_ reached a value of 0.8-1. During induction, CoCl_2_ (final concentration 0.2 mM) and L-arabinose (final concentration 2 g.L^-1^) were added, and incubation temperature was decreased to 23°C for 20 hours. Cultures were stopped and cells harvested by centrifugation (4400 g, 10 °C, 20 min), the pellet was resuspended in *lysis buffer* (50 mM HEPES pH 8.0, 150 mM NaCl, 0.2 mM CoCl_2_, 0.25 mg.mL^-1^ lysozyme, 0.1 mM PMSF and 10 µg.mL^-1^ DNAseI) and stored at -80 °C. Cells were then lysed by sonication (3×30 seconds Qsonica, Q700; Amplitude 45). As a pre-purification step, the lysate was heated for 30 min at 70°C. Cell debris and precipitated host proteins were eliminated by centrifugation (14000 g, 10 °C, 15 min). *Sso*Pox and its variants were concentrated by ammonium sulfate precipitation (75 %) and resuspended in 8 ml of *activity buffer* (50 mM HEPES pH 8.0, 150 mM NaCl, 0.2 mM CoCl_2_). Ammonium sulfate was eliminated by a desalting column step (HiPrep 26/10 desalting, GE Healthcare; ÄKTA Avant). The sample was concentrated to 2-3 mL for separation on exclusion size chromatography (HiLoad 16/600 Superdex^TM^ 75pg, GE Healthcare; ÄKTA Avant). Finally, purity was determined by SDS-PAGE and protein concentration was measured with a NanoDrop 2000 spectrophotometer (Thermo Scientific).

### Kinetic Assays

#### Generalities

Catalytic parameters were measured at 25°C in triplicate in 96-well plates using a reaction volume of 200 µL and recorded by a microplate reader (Synergy HT, BioTek, USA) in a 6.2 mm path length cell and using the Gen5.1 software, as previously explained.^31^ Graph-Pad Prism 6 software was used to derive catalytic parameters by fitting the data to the *Michaelis-Menten (MM)* equation. When V_max_ could not be reached, catalytic efficiency was determined by fitting the linear part of MM plot to a linear regression. For phosphotriester analogues, low concentration of substrates was used (10 µM), *k*_*cat*_/*K*_*M*_ were estimated using *one-phase decay* non-linear regression in Graph-Pad Prism 6 software. Catalytic parameters are presented in **Table 1** and **S2**, all the curves used for kinetic analyses are presented in **Figure S3** and **S4**.

**Table 1.** Catalytic parameters of selected *Sso*Pox variants against insecticides, an ester and lactones.

| Substrate | *SsoPox* | $k_{cat}$ (s<sup>-1</sup>) | $K_M$ (μM) | $k_{cat}/K_M$ (M<sup>-1</sup>.s<sup>-1</sup>) | Ratio/wt |
| --- | --- | --- | --- | --- | --- |
| **Ethyl-paraoxon** | *wt*<sup>+</sup> | $1,26 \pm 0,13 \times 10^1$ | $2,42 \pm 0,37 \times 10^4$ | $5,19 \pm 0,95 \times 10^2$ | 1 |
| | IH1 | $4,00 \pm 0,30 \times 10^{-2}$ | $8,81 \pm 1,41 \times 10^2$ | $4,38 \pm 2,16 \times 10^1$ | 0,08 |
| | IG7 | $1,69 \pm 0,20$ | $8,93 \pm 2,12 \times 10^2$ | $1,89 \pm 0,94 \times 10^3$ | 3,64 |
| | IIIC1<sup>#</sup> | $6,99 \pm 0,25 \times 10^1$ | $6,10 \pm 0,50 \times 10^2$ | $1,14 \pm 0,50 \times 10^5$ | 219,65 |
| | IIIB8 | $3,30 \pm 0,20 \times 10^{-1}$ | $5,05 \pm 0,68 \times 10^2$ | $6,47 \pm 2,62 \times 10^2$ | 1,25 |
| | IIIE11 | $3,00 \pm 0,20 \times 10^{-2}$ | $6,01 \pm 0,91 \times 10^2$ | $4,47 \pm 1,93 \times 10^1$ | 0,09 |
| | IVE2 | $3,30 \pm 0,20 \times 10^{-1}$ | $1,19 \pm 0,15 \times 10^3$ | $2,78 \pm 1,51 \times 10^2$ | 0,54 |
| | IVG2 | $2,20 \pm 0,20 \times 10^{-1}$ | $1,24 \pm 0,16 \times 10^3$ | $1,75 \pm 0,96 \times 10^2$ | 0,34 |
| | IVA4 | * | * | $1,06 \pm 0,02 \times 10^4$ | 20,42 |
| | IVE5 | * | * | $1,69 \pm 0,05 \times 10^3$ | 3,26 |
| | IVB10 | $3,06 \pm 0,11 \times 10^2$ | $1,11 \pm 0,78 \times 10^3$ | $2,75 \pm 1,46 \times 10^5$ | 529,87 |
| | IVE10 | $2,81 \pm 0,09$ | $2,04 \pm 0,24 \times 10^2$ | $1,38 \pm 0,39 \times 10^4$ | 26,59 |
| | VA7 | $2,40 \pm 0,10 \times 10^{-1}$ | $2,58 \pm 0,39 \times 10^2$ | $9,17 \pm 2,86 \times 10^2$ | 1,77 |
| | αsD6<sup>°</sup> | $3,22 \pm 0,18 \times 10^1$ | $1,09 \pm 0,12 \times 10^3$ | $2,95 \pm 1,55 \times 10^4$ | 56,84 |
| | W263I<sup>+</sup> | * | * | $1,21 \pm 0,06 \times 10^3$ | 2,33 |
| **Ethyl-parathion** | *wt*<sup>+</sup> | ND | ND | ND | ND |
|  | IH1 | ND | ND | ND | ND |
|  | IG7 | ND | ND | ND | ND |
| | IIIC1 | $8,20 \pm 0,40 \times 10^{-2}$ | $3,49 \pm 0,49 \times 10^2$ | $2,35 \pm 0,08 \times 10^2$ | ND |
|  | IIIB8 | ND | ND | ND | ND |
|  | IIIE11 | ND | ND | ND | ND |
| | IVE2 | $2,20 \pm 0,10 \times 10^{-3}$ | $5,10 \pm 0,84 \times 10^2$ | $4,44 \pm 1,81$ | ND |
| | IVG2 | $2,30 \pm 0,09 \times 10^{-3}$ | $3,00 \pm 0,35 \times 10^2$ | $7,75 \pm 2,57$ | ND |
| | IVA4 | $4,80 \pm 0,10 \times 10^{-1}$ | $3,92 \pm 0,25 \times 10^2$ | $1,22 \pm 0,45 \times 10^3$ | ND |
|  | IVE5 | ND | ND | ND | ND |
| | IVB10 | $2,86 \pm 0,19 \times 10^{-2}$ | $7,37 \pm 0,45 \times 10^2$ | $3,89 \pm 1,81 \times 10^1$ | ND |
| | IVE10 | $6,71 \pm 0,08 \times 10^{-3}$ | $1,45 \pm 0,17 \times 10^2$ | $4,63 \pm 1,15 \times 10^1$ | ND |
| | VA7 | $3,40 \pm 0,08 \times 10^{-3}$ | $9,52 \pm 1,04 \times 10^1$ | $3,63 \pm 0,77 \times 10^1$ | ND |
| | αsD6<sup>°</sup> | $7,35 \pm 0,36 \times 10^{-1}$ | $5,34 \pm 0,62 \times 10^2$ | $1,38 \pm 0,57 \times 10^3$ | ND |
|  | W263I | ND | ND | ND | ND |
| **pNP acetate** | *wt*<sup>°°</sup> | $1,70 \pm 0,07 \times 10^{-1}$ | $5,45 \pm 0,35 \times 10^3$ | $3,12 \pm 0,33 \times 10^1$ | 1 |
| | IH1 | $2,13 \pm 0,10 \times 10^{-2}$ | $5,80 \pm 0,67 \times 10^2$ | $3,67 \pm 1,56 \times 10^1$ | 1,18 |
| | IG7 | $7,08 \pm 0,99 \times 10^{-1}$ | $4,65 \pm 0,84 \times 10^3$ | $1,52 \pm 1,18 \times 10^2$ | 4,87 |
| | IIIC1 | $1,21 \pm 0,29 \times 10^{-2}$ | $7,12 \pm 2,03 \times 10^3$ | $1,71 \pm 1,41$ | 0,05 |
| | IIIB8 | $5,92 \pm 0,17 \times 10^{-2}$ | $1,86 \pm 0,09 \times 10^3$ | $3,18 \pm 1,97 \times 10^1$ | 1,02 |
| | IIIE11 | $2,28 \pm 0,07 \times 10^{-2}$ | $2,99 \pm 0,28 \times 10^2$ | $7,64 \pm 2,53 \times 10^1$ | 2,45 |
| | IVE2 | $2,70 \pm 0,04 \times 10^{-2}$ | $8,70 \pm 0,29 \times 10^2$ | $3,10 \pm 1,52 \times 10^1$ | 0,99 |
| | IVG2 | $4,41 \pm 1,13 \times 10^{-3}$ | $3,02 \pm 1,10 \times 10^3$ | $1,46 \pm 1,03$ | 0,05 |
|  | IVA4 | ND | ND | ND | ND |
|  | IVE5 | ND | ND | ND | ND |
| | IVB10 | $2,75 \pm 0,29 \times 10^{-2}$ | $2,11 \pm 0,36 \times 10^3$ | $1,31 \pm 0,81 \times 10^1$ | 0,42 |
| | IVE10 | $2,11 \pm 0,16 \times 10^{-3}$ | $1,63 \pm 0,21 \times 10^3$ | $1,29 \pm 0,77$ | 0,04 |
| | VA7 | $6,29 \pm 0,16 \times 10^{-2}$ | $2,02 \pm 0,08 \times 10^3$ | $3,11 \pm 1,97 \times 10^1$ | 1,00 |
| | αSd6 | * | * | $1,97 \pm 0,27 \times 10^1$ | 0,63 |
| | W263I | $6,71 \pm 0,27 \times 10^{-3}$ | $5,79 \pm 0,55 \times 10^2$ | $1,16 \pm 0,49 \times 10^1$ | 0,37 |
| **3-oxo-C10**<br>**AHL** | *wt*<sup>+</sup> | $4,52 \pm 0,10$ | $1,43 \pm 0,15 \times 10^2$ | $3,16 \pm 0,33 \times 10^4$ | 1 |
| | IH1 | $1,84 \pm 0,03 \times 10^{-2}$ | $6,79 \pm 0,91 \times 10^1$ | $2,71 \pm 0,36 \times 10^2$ | 0,01 |
| | IG7 | $2,70 \pm 0,13 \times 10^{-1}$ | $2,17 \pm 0,36 \times 10^2$ | $1,24 \pm 0,54 \times 10^3$ | 0,04 |
| | IIIC1 | $5,61 \pm 0,29 \times 10^{-1}$ | $5,72 \pm 0,81 \times 10^2$ | $9,80 \pm 3,58 \times 10^2$ | 0,03 |
| | IIIB8 | $1,64 \pm 0,06 \times 10^{-2}$ | $3,80 \pm 0,47 \times 10^2$ | $4,33 \pm 1,39 \times 10^1$ | 0,001 |
| | IIIE11 | $1,68 \pm 0,04 \times 10^{-2}$ | $6,43 \pm 1,29 \times 10^1$ | $2,61 \pm 0,34 \times 10^2$ | 0,01 |
| | IVE2 | $1,08 \pm 0,03 \times 10^{-2}$ | $2,10 \pm 0,26 \times 10^2$ | $5,14 \pm 1,23 \times 10^1$ | 0,002 |
|  | IVG2 | ND | ND | ND | ND |
|  | IVA4 | ND | ND | ND | ND |
|  | IVE5 | ND | ND | ND | ND |
| | IVB10 | $8,88 \pm 0,48 \times 10^{-3}$ | $4,19 \pm 0,72 \times 10^2$ | $2,11 \pm 0,67 \times 10^1$ | 0,001 |
|  | IVE10 | ND | ND | ND | ND |
| | VA7 | $2,60 \pm 0,10 \times 10^{-2}$ | $1,52 \pm 0,30 \times 10^2$ | $1,71 \pm 0,35 \times 10^2$ | 0,01 |
| | αSd6<sup>o</sup> | ND | ND | ND | ND |
| | W263I<sup>+</sup> | $6,00 \pm 0,90 \times 10^{-1}$ | $1,60 \pm 0,44 \times 10^3$ | $3,74 \pm 1,17 \times 10^2$ | 0,01 |
| **3-oxo-C12**<br>**AHL** | *wt*<sup>+</sup> | $1,01 \pm 0,13$ | $4,56 \pm 1,28 \times 10^2$ | $2,22 \pm 0,68 \times 10^3$ | 1 |
| | IH1 | $4,22 \pm 0,30 \times 10^{-3}$ | $1,79 \pm 0,42 \times 10^2$ | $2,36 \pm 0,72 \times 10^1$ | 0,01 |
| | IG7 | $6,63 \pm 0,51 \times 10^{-2}$ | $2,74 \pm 0,58 \times 10^2$ | $2,42 \pm 0,88 \times 10^2$ | 0,11 |
| | IIIC1 | $2,31 \pm 0,89$ | $2,17 \pm 1,11 \times 10^3$ | $1,06 \pm 0,80 \times 10^3$ | 0,48 |
|  | IIIB8 | ND | ND | ND | ND |
| | IIIE11 | $7,46 \pm 0,47 \times 10^{-3}$ | $1,27 \pm 0,33 \times 10^2$ | $5,88 \pm 1,44 \times 10^1$ | 0,03 |
| | IVE2 | $5,93 \pm 0,73 \times 10^{-2}$ | $7,50 \pm 1,70 \times 10^2$ | $7,91 \pm 4,29 \times 10^1$ | 0,04 |
|  | IVG2 | ND | ND | ND | ND |
|  | IVA4 | ND | ND | ND | ND |
|  | IVE5 | ND | ND | ND | ND |
|  | IVB10 | ND | ND | ND | ND |
|  | IVE10 | ND | ND | ND | ND |
|  | VA7 | ND | ND | ND | ND |
| | αSd6 | $1,65 \pm 0,21 \times 10^{-2}$ | $7,78 \pm 2,07 \times 10^2$ | $2,12 \pm 1,04 \times 10^1$ | 0,01 |
| | W263I<sup>+</sup> | $1,80 \pm 0,05$ | $1,78 \pm 0,49 \times 10^1$ | $1,01 \pm 0,28 \times 10^5$ | 45,50 |

#### Phosphotriesterase activity characterization

The kinetic assays were done in *activity buffer* (50 mM HEPES pH 8.0, 150 mM NaCl, 0.2 mM CoCl_2_). The conditions used for pesticides were as follows: ethyl-paraoxon, ethyl-parathion, and *p*-nitrophenol acetate hydrolysis took place at 405 nm (ε = 17000 M^-1^.cm^-1^) within a range of 0.05 to 2 mM as the substrate concentration, each solution made from a 100 mM stock solution in ethanol. For phosphotriester analogues, measurements were taken at 10 µM from a 100 mM stock solution in DMSO. We assumed that *K*_*M*_ ≫ [*S*], thus enabling us to estimate *k*_*cat*_/*K*_*M*_. Hydrolysis was measured by fluorescence (360/40, 460/40 nm).

#### Lactonase activity characterization

Lactonase kinetics were performed as described previously^34^. Kinetic parameters were determined in *lac buffer* (2.5 mM Bicine pH 8.3, 150 mM NaCl, 0.2 mM CoCl_2_, 0.25 mM Cresol purple and 0.5 % DMSO) over a concentration range of 0-2 mM for 3-oxo-C10 AHLs and 3-oxo-C12 AHLs. Hydrolysis was followed through measurement of absorbance at 577 nm (ε = 2923 M^-1^.cm^-1^).

### Production and Partial Purification of rHAChE

As previously reported, rHAChE was produced in *E. coli* Origami B cells transformed with pET32-rAChE-3G4 plasmid kindly provided by Moshe Goldsmith from the Weizmann Institute (Rehovot, Israel).^16^ Starter was realized in 2^x^YT medium (Trypton 16 g.L^-1^, yeast extract 10 g.L^-1^ and NaCl 5 g.L^-1^) supplemented with ampicillin (100 µg.mL^-1^) and grown overnight at 37°C and under stirring. Inoculates were added (1:50 dilution) to 3L of 2xYT medium with ampicillin and grown at 37 °C until OD_600 nm_ reach 1. Then IPTG was added (final concentration of 0.2 mM), and temperature was changed to 16°C. Cultures were grown for 24h, cells were harvested by centrifugation (4400 g, 4 °C, 20 min), resuspended with 300 mL of *rHAChE lysis buffer* (13 mM Tris pH 8.0, 33 mM NaCl, 10 mM EDTA, 10 % glycerol, 0.25 mg.mL^-1^ lysozyme) and stored at -80 °C. Cells were sonicated (3×30 seconds Qsonica, Q700; Amplitude 40). Cell debris were eliminated by centrifugation (14000 g, 10 °C, 15 min), and 0.1% of octyl glucoside was added. To eliminate more cell debris and host proteins a first ammonium sulfate precipitation was realized, increasing ammonium sulfate concentration from 0 to 40% followed by mixing for 2h at 4°C. As previously, a centrifugation was done (14000 g, 10 °C, 15 min), and a second ammonium sulfate cut was performed in the supernatant, increasing the concentration of ammonium sulfate from 40% to 50% followed by overnight precipitation of rHAChE at 4° C. A centrifugation was done (14000 g, 10 °C, 15 min) and the pellet was resuspended in *rHAChE buffer* (13 mM Tris pH 8.0, 33 mM NaCl, 10 mM EDTA, 10 % glycerol) with 0.1% of octyl glucoside. Ammonium sulfate was removed by injection on a desalting column (HiPrep 26/10 desalting, GE Healthcare; ÄKTA Avant). The sample was concentrated before separation on exclusion size chromatography (HiLoad 16/600 Superdex^TM^ 75pg, GE Healthcare; ÄKTA Avant). Activity of fractions containing partially purified rHAChE was measured using Ellman’s reagent (DTNB, 4 mM) and acetylthiocholine (2.5 mM), following the reaction at 412 nm over 10 min.^42^ Fractions with activity were gathered and concentrated to reach around 2-3 U.mL^-1^.

### rHAChE Inhibition Assay

Inhibition assays were performed as previously described.^16, 43^ Briefly, the assay was divided in two steps. First, a reaction mixture containing *Sso*Pox (at 0.1, 1 or 10 µM) and paraoxon (1.5 µM) was started in *activity buffer*, and samples were taken every minute between 0 and 10 min with a final sample at 20 min. The samples were incubated with rHAChE for 15 min. The second step consisted of measuring the remaining activity of rHAChE using Ellman’s reagent (DTNB, 4 mM) and acetylthiocholine (2.5 mM) as described above in *activity buffer* supplemented with DTNB 4 mM. The slopes were used to calculate the remaining percentage of rHAChE activity. Curves were fitted with *one-phase decay* equation using GraphPad Prism 6 software.

### Melting Temperature Determination

Melting temperatures (T_m_) were obtained using differential scanning fluorimetry (DSF).^44^ Experiments were performed on CFX96 Touch™ Real-Time PCR Detection System (Bio-Rad). *Sso*Pox variants were used at 1 mg.mL^-1^ in *Tris buffer* (50 mM Tris-HCl, pH 7) and with 1X SYPRO^®^ orange (Sigma-Aldrich). Denaturation was followed using FRET channel. Experiments were performed with temperature increasing from 35°C up to 95°C (at a rate of 0.5°C/15 sec). For the most thermostable variants guanidinium chloride or urea was used with a range between 0.5 and 3 M. Data were fitted with *Boltzmann sigmoidal* equation using GraphPad Prism 6 software (San Diego, California USA, www.graphpad.com), and T_m_ was extrapolated by *linear regression*.

### Crystallization

Crystallization assays were performed as previously described,^45^ using the hanging drop vapor diffusion method in 24-well VDX plates with sealant (Hampton research, California, USA). Different protein:precipitant ratios were tried (1:1; 1:2 and 1:3) and the plate were incubated at 292K. Crystals appeared after three days in a solution containing 25% (w/v) PEG 8000 and Tris-HCl buffer (pH 8.5). To obtain quality crystals, micro seeding was performed. Protein crystals were harvested from a drop, placed in 5 µL of mother solution and vortexed for 1 min. Several dilutions of seeds were performed (50, 75 and 100-fold) and 0.1 µL of these solutions were added to drops containing different protein:precipitant ratios. Attempts were made to co-crystallize these variants with Diethyl-4- methylbenzylphosphonate and crystals were produced for the variants IVA4, IG7 and IVB10 but the compound could not be modelled.

### Data Collection and Structure Refinement

Crystals were transferred into a cryoprotectant solution composed of 20 % (v/v) glycerol and crystallization solution. Crystals were then flash-cooled in liquid nitrogen. X-ray diffraction data were collected at 100 K using synchrotron radiation at the 23ID-B beam line (APS, Argonne, USA). X-ray diffraction data were integrated and scaled with the *XDS* package **(Table S3** and **S4)**.^46^ The phases were obtained performing a molecular replacement with MOLREP or PHASER using the structure of *Sso*Pox-*wt* (PDB ID 2VC5) as model.^47, 48^ The models were built with *Coot* and refined using REFMAC^49, 50^ and PHENIX^51^. Structures were validated using Molprobity^52^. Structure illustrations were obtained using PyMOL.^48^

### Structural Analysis

Active site cavities were determined using CAVER Analyst 2.0 beta^53^ and a 1.4A *probe* and *large probe* 3.00. For IIIC1, these settings resulted in artifactually small multiple cavities. Therefore, calculations were done by allowing a probe up to 4.00, and the program chose a max probe of 3.05 for this variant. Average B-factors were calculated using *Baverage* from the CCP4 program suite v.8.0.004.^54^

## 3. RESULTS AND DISCUSSION

### Combinatorial Library Screening Yields Higher Activity, Highly Promiscuous Mutants

Initial screening of the library using an ethyl-paraoxon colored colony assay allowed to identification of 430 active variants **(Figure S1)**. These variants were subsequently submitted to further screenings using nine substrates including organophosphorus compounds (ethyl-paraoxon and previously reported phosphotriester analogues^55, 56^), acyl-homoserine lactones (3-oxo-C10 AHL and 3- oxo-C12 AHL) and arylester (*para*-nitrophenyl acetate). The results from these screens are presented as a heat map in the supplementary materials **(Figure S2)**. From these results, 12 highly active variants were identified and subsequently submitted to complete kinetic characterization towards the nine previously listed substrates and ethyl parathion. The 12 highly active variants harbor 2 to 6 mutations **(Table S5)**. The obtained kinetic parameters can be compared to those of the wild-type enzyme (*Sso*Pox-*wt*) and two previously reported, improved lactonase and phosphotriesterase variants of *Sso*Pox (*i.e*. W263I and αsD6^34, 36^); (**Table 1 and S2, Figure 2, Figure S3 and S4)**. Data reveal that the variants IIIC1 and IVB10 exhibit 220- and 530-fold increase in catalytic efficiency with ethyl-paraoxon as a substrate as compared to *Sso*Pox-*wt*. In fact, variant IVB10 is 10 times more active than the highly active variant αsD6 obtained by our group^36^ and ∼5-fold more active than the *4mut* variant previously reported^40^. Other activity improvements were noted. For example, the variant IG7 shows a ∼5-fold increase in catalytic efficiency for the arylesterase activity (**Table 1**). Both arylesterase and phosphotriesterase were improved with a similar magnitude for IG7, as compared to *Sso*Pox-*wt*. Interestingly, large improvements of phosphotriesterase and arylesterase activity appear concomitant with a drastic reduction in their lactonase activity (**Figure 2**). These activities therefore appear to functionally tradeoff in the *Sso*Pox enzyme. Overall, several variants, such as IIIC1 and IVE2, appear significantly more promiscuous than *Sso*Pox-*wt* (**Figure 2**).

**Figure 2.**
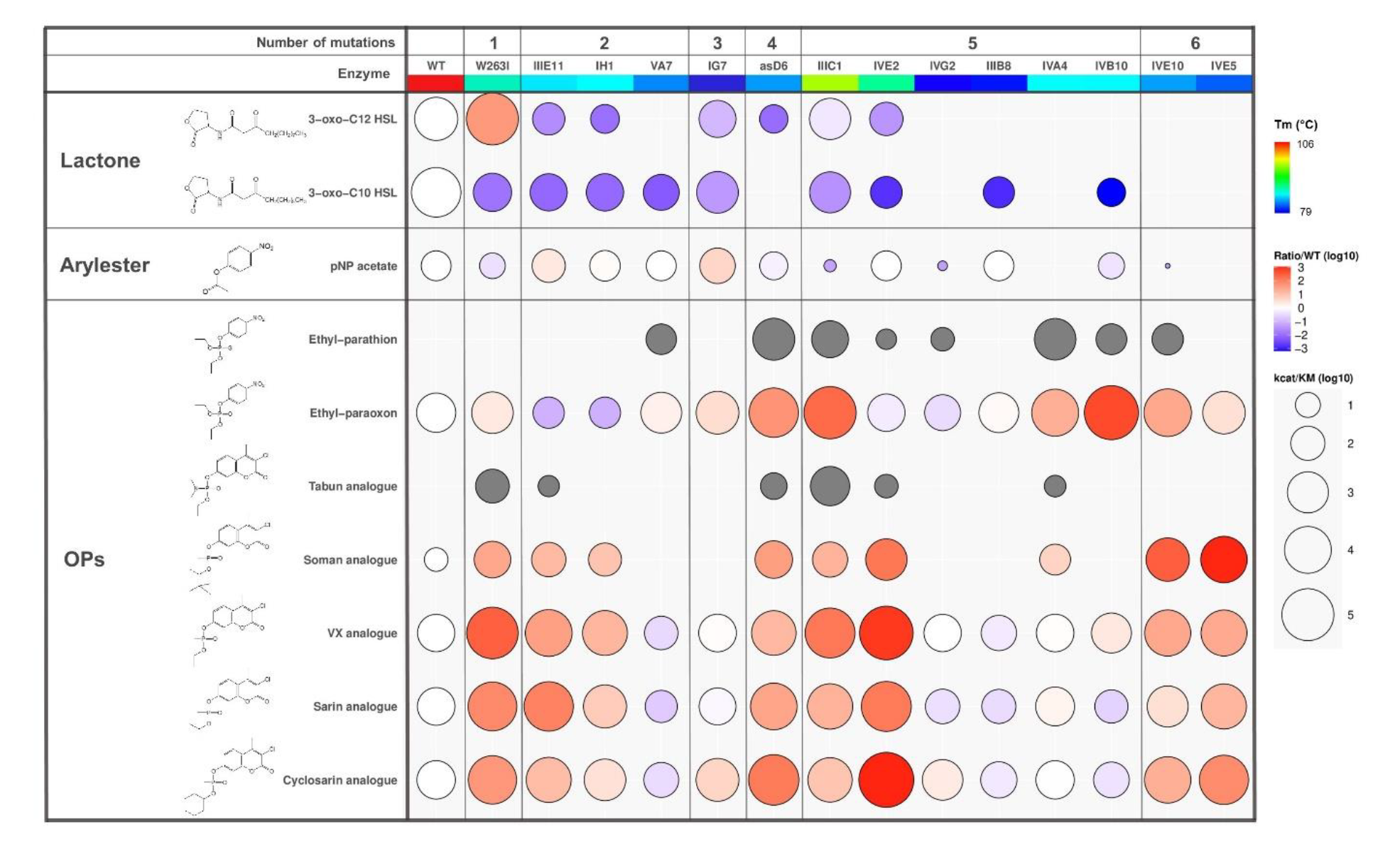
Enzymatic promiscuity and stability profile of *Sso*Pox variants. The heat map shows the selected variants catalytic efficiency values (circle size) and fold change (fill colors) compared to *Sso*Pox-wt, in logarithmic values. Grey fill indicates no calculated ratio relative to the WT due to lack of measured WT activity towards that substrate. Each column represents the data for a different *Sso*Pox variant while each row shows the results for a different substrate. Variants are organized by number of combined mutations (top row). The second table row indicates the variants names and their melting temperature (Tm) values (specific values are given in **Table S3**). The heat map was realized using the ggplot2 package on R.

### Improved Variants IIIC1 and IVA4 Do Not Show Any Thiono-effect

Remarkably, most (7 out of 12) characterized variants showed detectable ethyl-parathion activity, while *Sso*Pox-*wt* does not (**Table 1**). The absence of activity against ethyl-parathion is intriguing because it differs from ethyl-paraoxon only by the substitution of the terminal oxygen atom on the phosphorus atom into a sulfur atom (**Figure 2**). This behavior can be related to a thiono-effect: some PLLs were previously described to show marked preference for oxono-phosphotriesters,^57^ where PTEs do not show this pattern.^58^ The thiono-effect varies in the different variants: the paraoxonase/parathionase activity levels ratio are ranging from ∼9 to 7069 for variants IVA4 and IVB10, respectively. We note that all variants with measurable ethyl-parathion activity levels contain mutations at positions Y97 and Y99, including αsD6, generated in a previous study from our group.^36^ In particular, Y97 is relatively conserved in metallo-lactonases^12^ and was shown in other PLLs to be strongly charge_-_coupled to the metal cation β.^59, 60^ Interestingly, these tyrosine residues are not present in PTEs, enzymes that typically show very low thiono-effects. However, the mutation of Y97 or Y99 does not appear to be sufficient to reduce the thiono-effect, since some variants (e.g. IIIE11, IH1, IIIB8, IVE5) containing these mutations show no ethyl-parathion activity (**Table 1**).

### Some Variants Show Very Large Phosphotriesterase Activity Increase

The 12 selected variants were submitted to further kinetic characterization against several organophosphorus compounds. Specifically, coumaric analogues of sarin, cyclosarin, tabun, soman and VX were previously described.^62^ *Sso*Pox-*wt* exhibits low catalytic efficiency values against VX, sarin, cyclosarin and soman analogues (k_cat_/K_M_ values of 2.74 x 10^2^, 3.00 x 10^2^, 3.93 x 10^2^ M^-1^.s^-1^ and 7.63 M^-1^.s^-1^, respectively), and no detectable activity against the tabun analogue **(Table S2, Figure S4)**. Previous engineering efforts on *Sso*Pox lead to the improved variants W263I and αsD6 with increased lactonase and phosphotriesterase activities respectively.^34, 36^ We here measure these variants with new substrates, and show that the mutant W263I is 297.4-fold more active against the VX coumarin analogue, while the variant αsD6 showed a 125.4-fold increase against the cyclosarin analogue, as compared to *Sso*Pox-*wt*. In addition, both W263I and αsD6 show low, yet measurable activity for the hydrolysis of tabun analogue (**Figure 2, Table S2)**. The new variant IVE2 was found to be the most efficient variant against sarin, cyclosarin and VX analogues with k_cat_/K_M_ values reaching 3.90 x 10^4^, 4.38 x 10^5^ and 2.15 x 10^5^ M^-1^.s^-1^, respectively (**Figure 2, Table S2)**. This represents considerable improvements over *Sso*Pox-*wt*, specifically 130-, 1114- and 784-fold. When compared to W263I and αsD6, IVE2 shows higher activity for all tested analogues, and is ∼9-fold more active against the cylosarin analogue **(Table S2)**. Other variants are also largely improved: the variant IVE5 showed 1076-fold enhancement compared to *Sso*Pox-*wt* with the soman analogue as substrate. The variant IIIC1 showed improvement towards all five tested analogues and was the most active variant for the hydrolysis of the tabun analogue (k_cat_/K_M_ value of 5.28 x 10^2^ M^-1^.s^-1^).

Overall, a mutation at the position W263 is present in numerous improved variants against the coumarin analogues, including W263I, αsD6, IVE5, IVB10, IVE10, IIIC1 and IVE2. Other variants show improved phosphotriesterase activity without a mutation at position W263, but contain mutations at both the Y97 and Y99 positions, including IVA4, IVE2, IIIE11 and IH1. This may suggest that different strategies are at play to increase activity: mutating W263 mainly affects the loop 8 conformation^34^ and mutations at Y97 and Y99 sites may alter the cavity size, its electrostatic properties, and the potential for interactions with the substrate.

The best identified variants in degrading organophosphorus compounds (*i.e.* IIIC1, IVB10, IG7, IVE10 and IVA4) were subjected to further investigations. Specifically, we adapted a previously reported procedure based on the inhibition of Human AChE^16, 43^ to evaluate their capacity to protect the enzyme target of organophosphorous compounds. We used a previously reported stabilized recombinant variant of Human AChE (rHAChE).^16, 63^ rHAChE inhibition assays were performed with ethyl-paraoxon, although rHAChE inhibition by this substrate is not irreversible. rHAChE activity was measured at different time points to evaluate the rate of detoxification of the phosphotriester by the *Sso*Pox variants. The results show that IIIC1 and IVB10 were the fastest mutants to degrade the ethyl-paraoxon and thereby protect rHAChE from inactivation. The two mutants, used at a concentration of 0.1 µM, degraded the phosphotriester in 3 minutes, whereas *Sso*Pox-*wt,* tested at a 100-fold higher enzyme concentration, required more than ten minutes **(Figure S5)**. These results were in accordance with the much higher observed catalytic efficiency values for the variants IIIC1 and IVB10. Variant IIIC1 was recently evaluated for *in vivo* detoxification of organophosphate and results showed it could protect mice from ethyl-paraoxon.^38^

### Biochemical Properties of the Improved Variants

The biochemical properties of the obtained mutants, namely their kinetic properties and their thermal stability, were grouped in a heat map for visualization and analysis (**Figure 2**). This was performed by compiling a heat map of the overall activities and also by clustering the substrates by type and the variants according to the number and the type of mutations. The melting temperatures were determined for the selected variants **(Table S5)**. They show that all variants have reduced thermal stability as compared to *Sso*Pox-*wt*. In fact, the most stable mutant, IIIC1, shows a T_m_ value of 96.3°C, a 10°C reduction from the *Sso*Pox-*wt* T_m_ value (106°C). The least active mutant exhibits a T_m_ value of 79.1°C, a nearly 27°C drop. This is evidence that the identified mutations and / or combination of mutations are destabilizing. Yet, all the mutants remain considerably thermostable, with T_m_ values above 79°C **(Table S5)**.

Analysis of the kinetic properties of the variants reveals that while their phosphotriesterase activity was improved, their lactonase activity was greatly decreased. In some variants, some key active site residues are mutated. For example, Y97 and Y99 are mutated in the variants IIIE11 and IH1. Y97 in particular was previously reported to be involved in the lactone ring positioning in *Sso*Pox, yet it does not appear required for the lactonase activity.^45^ Other observations can be made from the kinetic parameters analysis. It seems that the Y97F mutation is present in mutants with the largest activity increase for ethyl-paraoxon (*i.e.* IIIC1, IVB10, IVE10). This is consistent with previous description of the narrowness of the binding cleft in the catalytic subsite of *Sso*Pox-*wt*, partly due to Y97, that prevents proper binding of the bulkier phosphotriester molecules.^31^

### The Selected Mutations Alter the Active Site Loops Conformation and Flexibility

The X-ray structures of some of the most improved variants, namely IG7, IIIC1, IVA4, IVB10 and IVE2 could be resolved **(Table S3)**. Additional structures for IG7, IVA4 and IVB10 derived from co-crystallization trials with a non-hydrolysable paraoxon analogue have failed to capture a bound state but allowed capture of alternative, highly disordered conformations **(Table S4)**. For some variants, molecules were bound to the active site, as evidenced by the examination of the electron density maps. Because the observed density appeared tetrahedral, a phosphate anion was modelled in the structures of the variants IG7, IVA4, IVB10. Yet, density maps suggest it might correspond to a larger, unidentified molecule (**Figure S6** for variant IVB10). The molecules bind onto and between the two metal cations, in a similar fashion in all structures and similarly to a previous observation made in the structure of another lactonase.^64^

Overall, the obtained structures are similar to that of *Sso*Pox-*wt*. Indeed, the r.m.s.d. on all atoms between *Sso*Pox-*wt* and IVE2, IG7, IIIC1, IVA4 and IVB10 is 0.29Å, 0.30Å, 0.29Å, 0.31Å and 0.30Å, respectively. Interestingly, consistent with observations of previous mutants,^34^ most of the changes between the structures occur in loops 7 and 8. This may be due to the localization of the mutations in the protein structure, being mostly situated in the active site vicinity **(Figure S7)**. The largest changes are related to the loop 8 conformation (**Figure 3**). In fact, it appears that there are two groups of loop 8 conformation: a conformation that is similar to that of *Sso*Pox-*wt* that is adopted in the variant IIIC1 (**Figure 3**), and a second conformation, extended, that is present in the structures of variants αsA6^36^, IVE2, IG7, IVA4 and IVB10 (**Figure 3**). Interestingly, the variant IIIC1 is also the crystallized variant with the highest T_m_ value **(Table S5)**, and consistently, the thermal B-factor for loops 7 and 8 is lower than that of other variants (**Figure 3**). In regards to loop 7, the variants show similar conformation to that of *Sso*Pox-*wt* with the exception of IVB10 (**Figure 3**).

**Figure 3.**
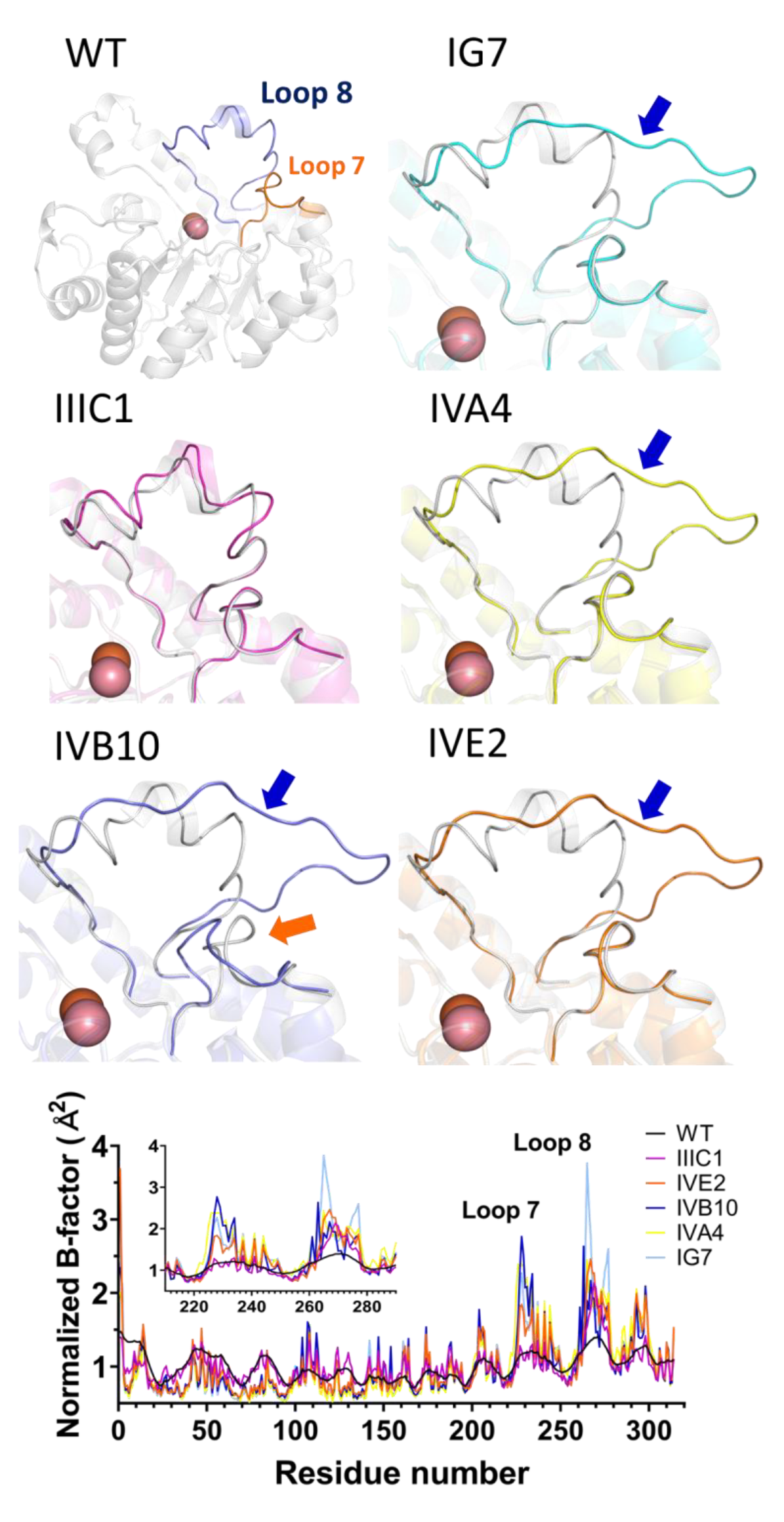
The improved variants show different active site loops 7 and 8 conformations. Active site regions of the variant structures are superimposed with the *Sso*Pox-*wt* structure (light grey; PDB ID 2VC5). The active site location is indicated by the presence of the bi-metallic catalytic center shown as spheres. The *Sso*Pox-*wt* structure and its loops 7 and 8 highlighted. Significant loop conformation changes are indicated with arrows (blue for loop 8; orange for loop 7). *Bottom graph*: Normalized thermal B-factor values of the different structures as a function of residue number. A zoomed inset for the loop 7 and 8 sequence region is shown.

The active site loops 7 and 8 not only adopt different conformations but also show different conformational flexibility. The structures highlight that the selected mutations contribute to this increased flexibility of the loops. In fact, for the additional structures solved for the variants IG7, IVA4 and IVB10, much of loops 7 and 8 could not be modelled due to the lack of electronic density, suggesting high levels of disorder **(Figure S8)**. This is confirmed by the alternative structure of the variant IG7 that shows low metal occupancy (modelled at 0.5 occupancy) and a possibly partially decarboxylated lysine (KCX137). Similar observations could be made for the alternative structure of the variant IVA4, albeit to a lesser extent. The latter structure further illustrates this increased conformational flexibility: it captures non-productive enzyme conformation caused by loop disorder **(Figure S9A)**. In this conformation, a fragment of loop 8 is bound to the bi-metallic active site center and thereby is a non-productive enzyme conformation **(Figure S9B)**. Residue C258 interacts with the cobalt cation and C259 forms a disulfide bridge with the mutated position L72C. Other structures reveal partial loop fragments that could be modelled **(Figure S8)**; but the structures show overall low electronic density to support robust loop modelling. For the more ordered structures with discrete loop conformations, the electron density supports model building for the active site loops **(Figure S10)**. The analysis of the normalized thermal motion B-factor confirms the increased flexibility of the loops 7 and 8 (**Figure 3** and **Figure S11)** as compared to *Sso*Pox-*wt* (PDB ID 2VC5). This higher mobility is consistent with the observation that all of the improved variants are destabilized compared to *Sso*Pox-*wt*, as shown by their reduced melting temperature values **(Table S5)**.

The changes in the enzyme’s active site for the conformation of loops 7 and 8 leads to a change in the substrate binding cavity size and shape. In fact, all of the variants’ structure showed significantly reshaped active site cavities **(Figure S12)**. In addition to the reshaping of the active site due to loop conformation changes, we note that numerous selected mutations include mutations into residues with smaller side chains that increase the volume of the binding cleft in the vicinity of the bimetallic center. This is true for the mutations V27G (IG7 and IVA4), V27A (IVB10), Y97F (IIIC1 and IVB10), Y97I (IVA4), Y97L (IVE2), and Y99F (IIIC1, IVA4 and IVB10). This local enlargement around the bi-metallic active site center may allow accommodation of the phosphoester groups that are bulkier than the lactone ring. However, the overall cavity size of the variants did not increase. In fact, each are smaller than that of *Sso*Pox-*wt*, and this is consistent with the observed large decreases in the lactonase activity for these variants, because AHLs can be much longer molecules that common phosphotriesters. The cavity volumes are 704.5, 519.0, 523.4, 382.5, 358.8 and 321.6Å^3^ for *Sso*Pox-*wt*, IVA4, IIIC1, IG7, IVB10, and IVE2, respectively. We note that the variant IIIC1, with similar loop conformation to *Sso*Pox-*wt*, retains a larger cavity size and high levels of lactonase activity. With the exception of variant IVA4, variants with smaller cavities show the lowest lactonase activity levels. These observations suggests that the size of the active site cavity contributes to modulating the activity profile of SsoPox.

### Changes in Interaction Networks and Alteration of the Active Site Loop Conformation

To better understand the molecular determinants for the observed conformational changes, we analyzed the effects of the mutations on the interaction network between the loops and the structure core (**Figure 4**). The analysis of the structure of the variant IG7 suggests a key role for mutation I280T. This residue anchors the C-termini of the active site loop 8 into the protein hydrophobic core. The mutation I280T may decrease this anchorage, and the threonine residue creates new interactions with E52 main and side chain, via two discrete conformations. This rearrangement causes the side chain of S279 to interact more closely with the side chain of T281 (3.2Å in IG7, 3.4Å in the *Sso*Pox-*wt* structure) and repositions the side chain of E52.

**Figure 4.**
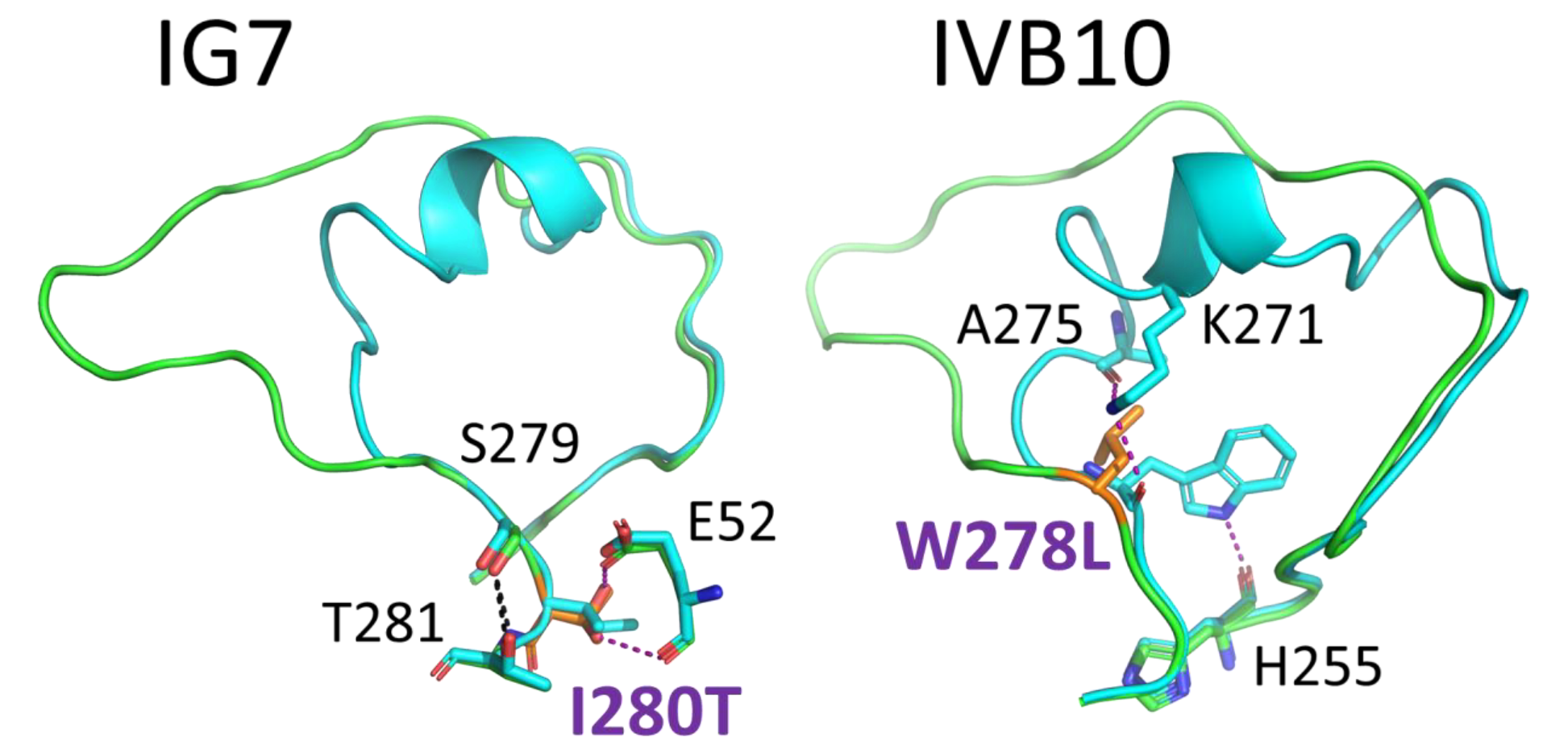
Mutations contribute to active site loops conformational changes. The variants structures (in green) are superposed onto the structure of *Sso*Pox-wt (in cyan). Mutated residues are highlighted in orange. Interactions are shown as dashed lines, and interactions of interest are shown as purple dashed lines.

For some variants, the structure does not immediately reveal a possible mechanism for the altered conformation. For example, the variant IIIC1 that also possesses I280T shows very similar features: a new interaction between I280T and E52 too and S279 interacting more closely with T281. Yet, in this case, these changes are not sufficient to alter the conformation of the loop 8 that adopts a *wt*-like conformation in the resolved IIIC1 structure **(Figure S13)**. Similarly, the mutations present in the IVA4 and IVE2 variants are not directly connected to the observed conformational changes. For these mutants, alteration of the active site loops conformation may originate from the destabilizing nature of the selected mutations, rather than a specific mechanism.

The structure of variant IVB10 suggests that the mutations cause the disruption of an intricate network of hydrogen bonds that exist in the *wt* structure. This disruption results in the more open loop 8 conformation. In the *wt* structure, the side chain of W278 forms a hydrogen bond with the main chain of H255 and its main chain interacts with K271 and A275. The mutation at W278L of variant IVB10 completely disrupts this interaction network, and an open loop conformation is observed (**Figure 4**).

Other factors are guiding the changes in loop 8 conformation. In particular, a difference between the variant IG7 and IIIC1 is the nature of the residue in position 263, W (IG7) or V (IIIC1), that was previously shown to affect the loop 8 flexibility and conformation.^34^

### Selected Mutations Modulate Conformational Sampling

Four of the variants could be solved at resolutions at which anisotropic refinement could be performed and improved the maps quality (IG7, IVE2, IVB10, IVA4). The available high-resolution data confirms the high structural mobility of the loops as illustrated by the corresponding electron density **(Figure S10).** Examination of the anisotropic atomic displacement parameters of the active site loops 7 and 8 reveals subtle differences between the variants (**Figure 5**). First, for most variants, the loop disorder is directional (**Figure 5**). Interestingly, with some variants, the intensity of the directionality and the direction itself can differ. For instance, variants IG7 and IVE2 exhibit a loop 8 with highly directional anisotropic thermal ellipsoids. In contrast, variants IVB10 and IVA4 exhibit lower degrees of directionality for both loop 7 and 8. These observations seems to define conformational clusters: despite having adopted similar conformation, the loop sampling may be different in these different variants. The notion that fine sampling can be captured in crystal structure has been recently exploited to unravel mechanisms^65, 66^. Different conformational clusters were previously described in PTEs^14, 67^ and hypothesized in *Sso*Pox^34^. Interestingly, these distinct structural clusters also correspond to functional clusters: variants IG7 and IVE2 both show high lactonase and esterase activity, while variants IVB10 and IVA4 show very low lactonase and esterase activity.

**Figure 5.**
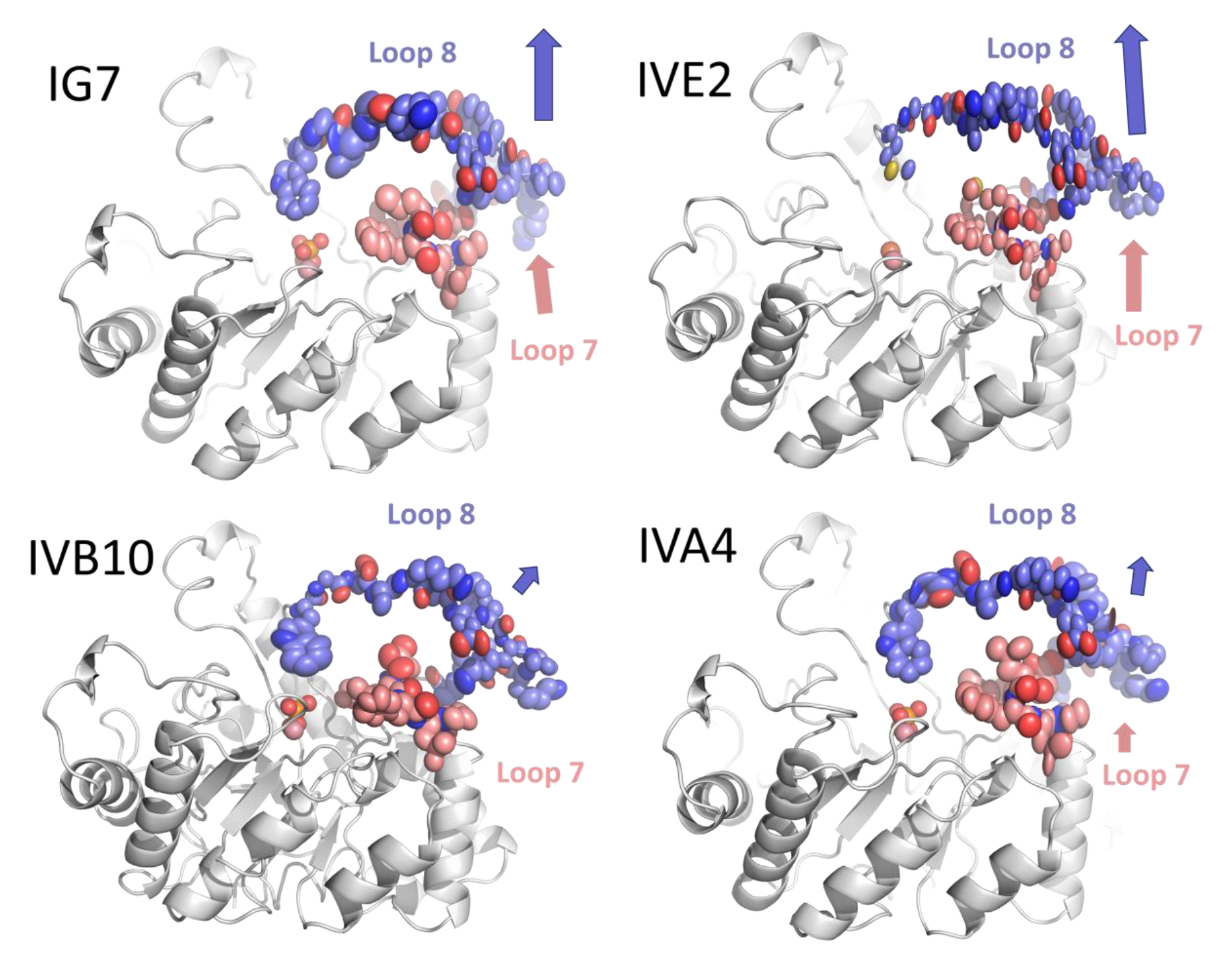
Active site loop conformation and anisotropic thermal B-factors ellipsoids. Arrows are an approximate representation of the direction and magnitude of the active site loops’ anisotropy. Metal cations and the bound phosphate anion are shown as spheres. The analysis was performed using PyMOL.

The importance of conformational dynamics in enzyme catalysis has been documented but remains a hot topic of discussion^68–71^. Yet, conformational dynamics have emerged as a key player to explain the creation of new enzymatic activities ^14, 69^. The results presented in this study suggest that conformational changes may be necessary but might not be sufficient for changing an enzyme’s activity. This is evidenced by the considerable remodeling of *Sso*Pox’s binding cleft, directly caused by the conformational changes of the active site loops 7 and 8. However, while these changes appear important, they may not be sufficient: IIIC1 is a mutant with improved phosphotriesterase activity and shows loop conformations similar to *Sso*Pox-wt. Examination of the high-resolution diffraction data and the anisotropic atomic displacement parameters for the active site loops suggest that conformational sampling, and the directionality of the latter is related to the enzyme’s activity profile. We surmise that loop sampling direction could favor or disfavor productive enzyme association with specific substrates. Such possible importance of conformational directionality was previously highlighted in the swiveling motion between its central domain and its nucleotide binding domain^72^, as well as loop dynamics in adenylate kinase^73^. Such subtle changes constitute an additional informational level beyond the existence of discrete conformations, and could improve the understanding of the contributions from distant mutations to catalysis and activity profiles.

## 4. CONCLUSION

In this work, we probed the possibility of redesigning the active site of the lactonase *Sso*Pox using structure-based design and combinatorial libraries. We obtained mutants with largely improved catalytic abilities against phosphotriesters, and generally more promiscuous. The best variants are 1,000-fold better compared to the wild-type enzyme. Remarkably, these improvements came at the cost of the initial lactonase activity: some of these variants are now specialized phosphotriesterases with specificity shifts in the vicinity of ∼1,000,000-fold. This is in contrast with previous approaches on the evolution of phosphotriesterase enzymes (PTEs), where bi-functional enzymes with relatively high proficiency could be obtained using random mutagenesis before significant tradeoffs was observed^74^. In this work, it may have been caused by the active site shape redesign approach, and the collected structural data informs about the active site requirements for both the lactonase and the phosphotriesterase activity. The designed mutations and their combinations appear to be destabilizing and considerably reshape the enzyme active site cavity via the reorganization of the active site loops. Changes in the loops correlate with the gain in promiscuous phosphotriesterase activity. Notably, high resolution structural data and analysis of the anisotropic atomic displacement parameters suggests loop conformational sampling and its directionality that seem to correlate with accepted substrates. While these observations appear critical to future engineering projects on SsoPox to improve its phosphotriesterase or lactonase activities, they also hint at the possible importance of loop sampling direction in defining enzymes’ activity profiles.

## ACKNOWLEDGEMENT

We are very grateful to Dr. Moshe Goldsmith from the Weizmann Institute (Rehovot, Israel) for kindly providing the rHAChE plasmid. We are very grateful to the scientists at the Advanced Photon Source (APS, Argonne, IL, USA) and particularly the beamline scientists and coordinators at 23ID-B for their assistance.

