## Supplementary material for "Changes in Active Site Loop Conformation Relate to the Transition toward a Novel Enzymatic Activity"

[#]Correspondence:

### Summary

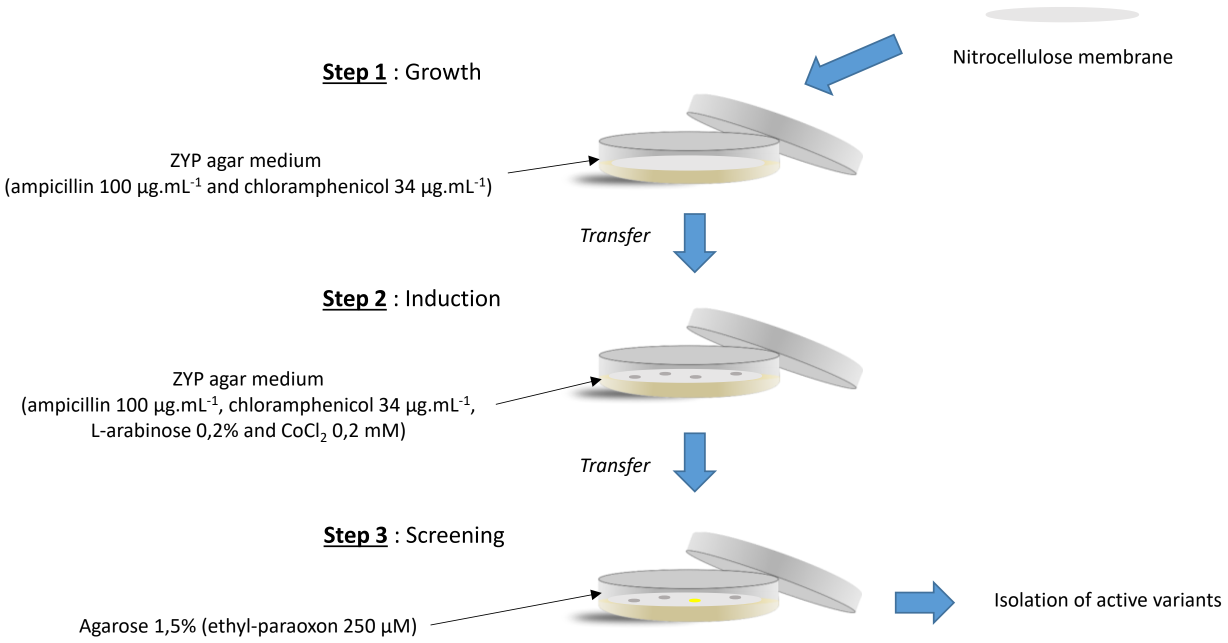

**Figure S1. Screening procedure.**
Description of color-based solid screening procedure of *Sso*Pox variants retaining activity on ethyl-paraoxon (250 µM) using nitrocellulose membranes.

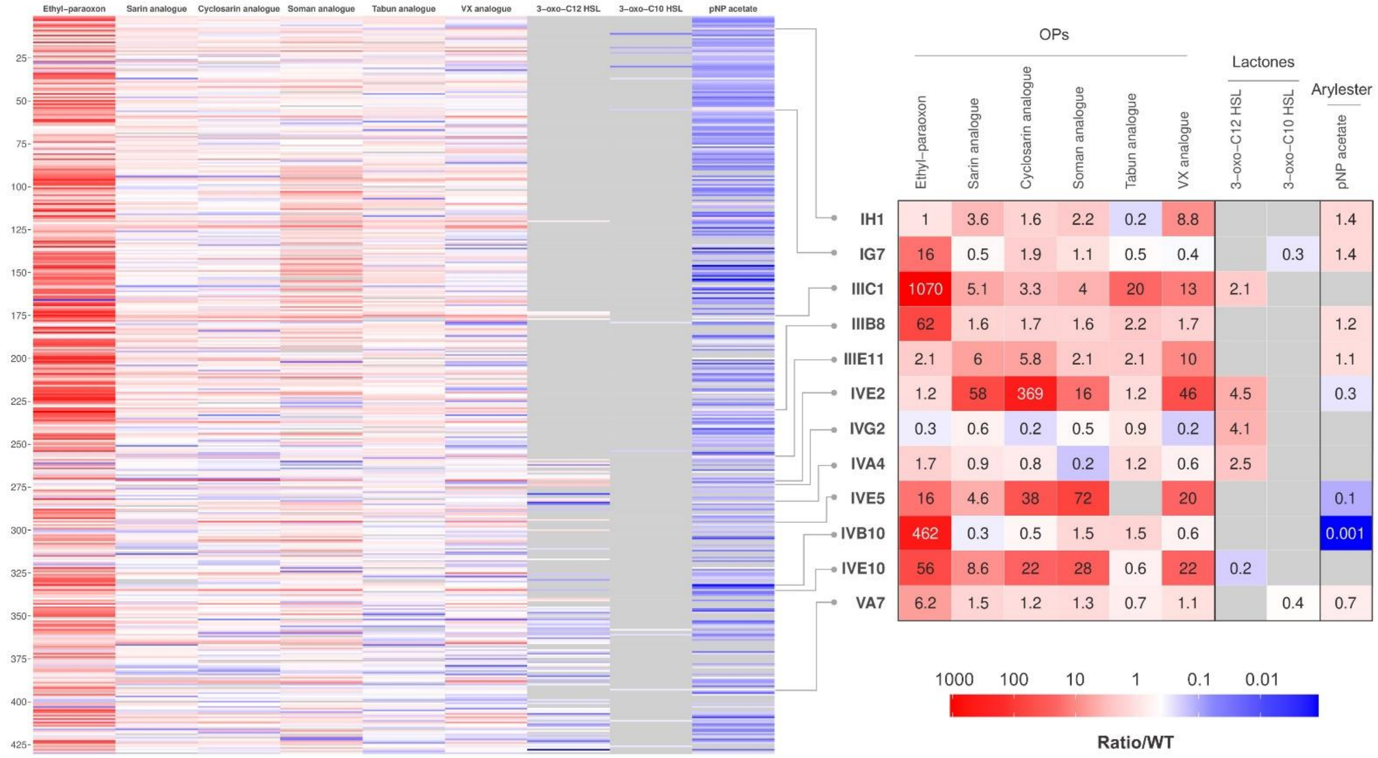

**Figure S2. Screening results.**

Screening results of the 430 *Sso*Pox variants assayed with 9 substrates as compared to wild-type enzyme. The heat map on the left displays the activity ratio of all 430 tested variants towards 9 substrates as compared to the wild type enzyme. Gray corresponds to no activity detected. Variants selected for further studies are identified with gray linkers and their screening activity ratios for each substrate are shown in the heat map on the right. Heat maps were realized using the ggplot2 package on R.

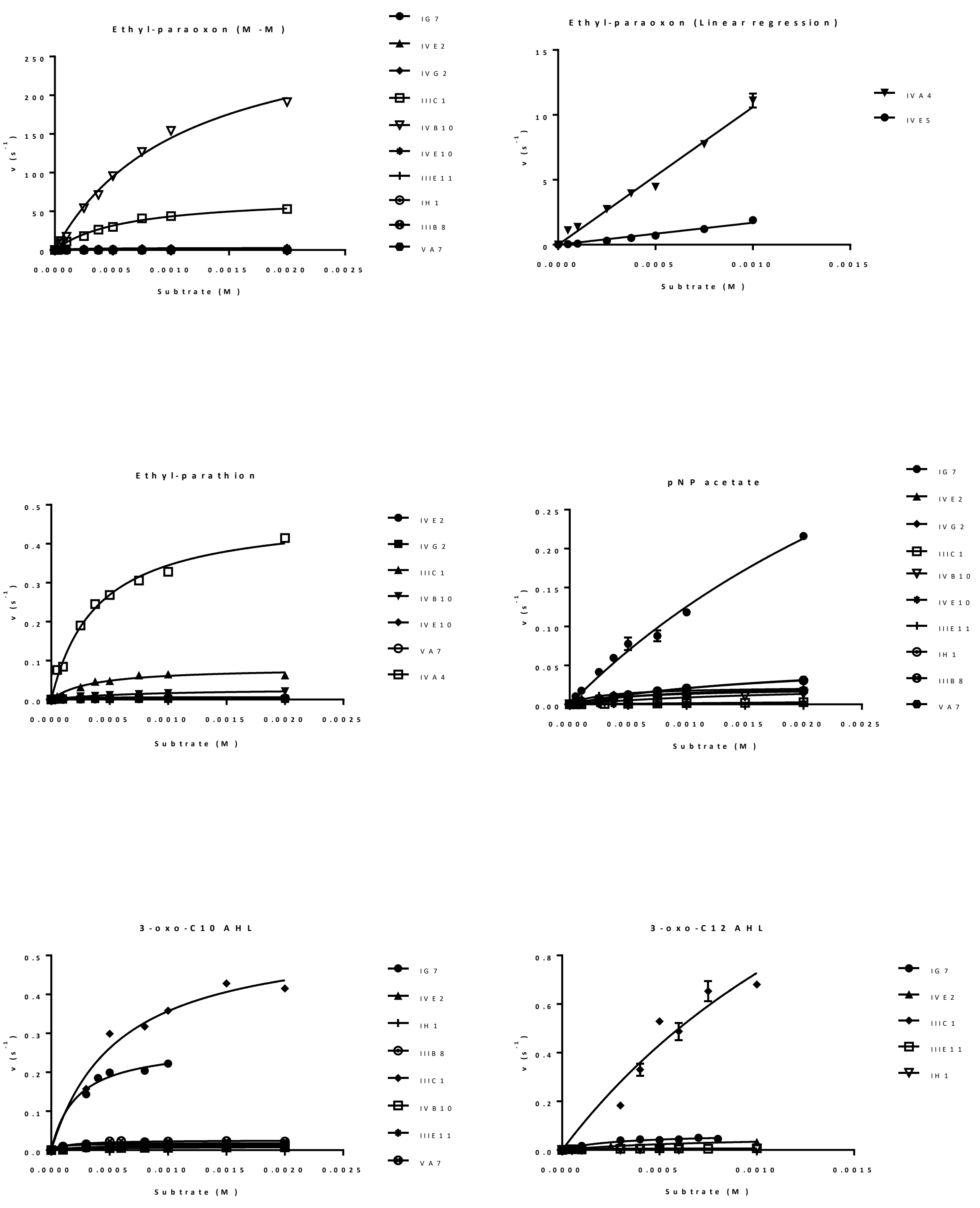

**Figure S3. Kinetic parameters determination.**

Catalytic parameters curves of *Sso*Pox variants on ethyl-paraoxon, ethyl-parathion, pNP acetate, 3-oxo-C10 AHL and 3-oxo-C12 AHL.

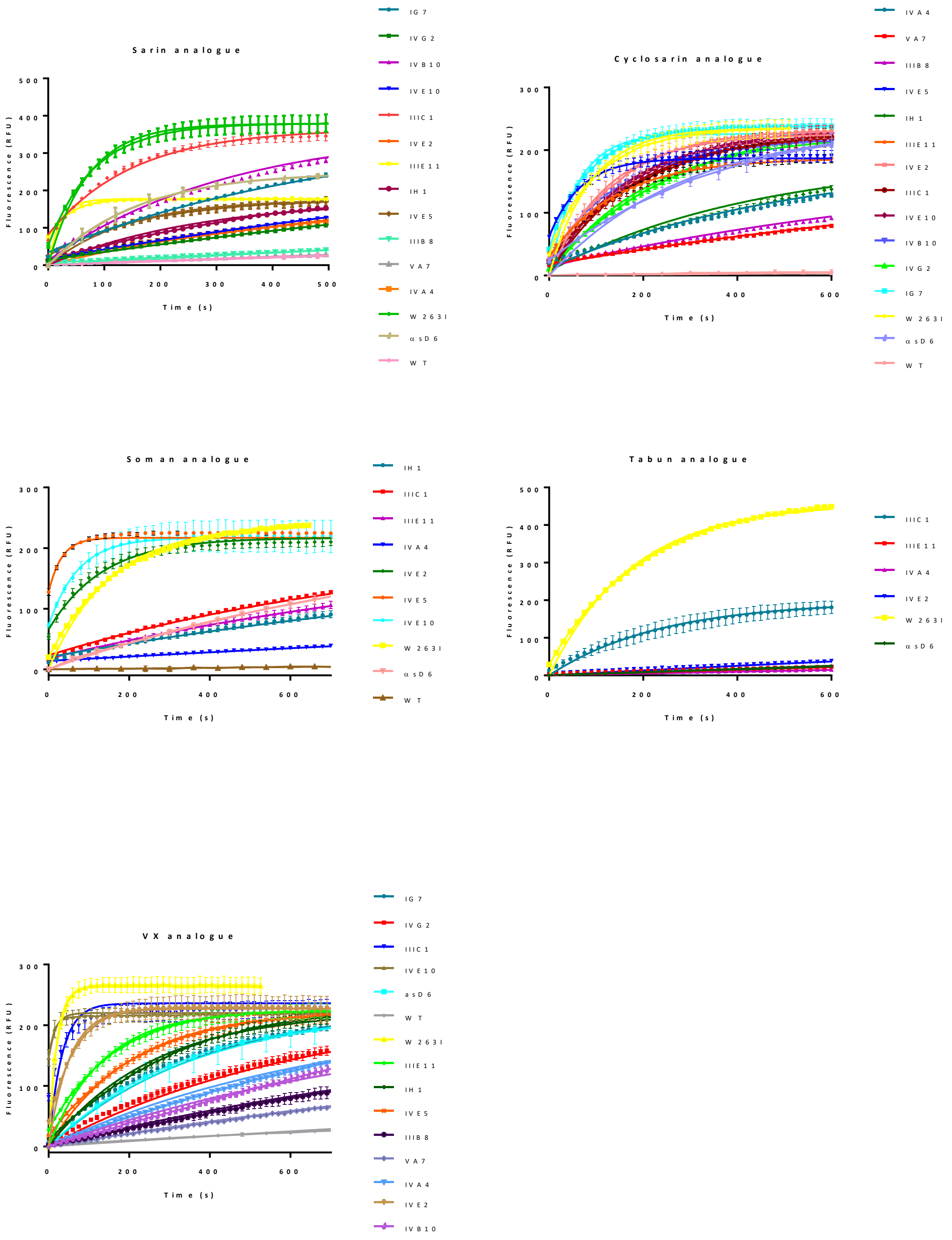

**Figure S4. Kinetic parameters determination for phosphotriester analogues**.

Catalytic parameters curves of *Sso*Pox variants on sarin analogue, cyclosarin analogue, soman analogue, tabun analogue and VX analogue.

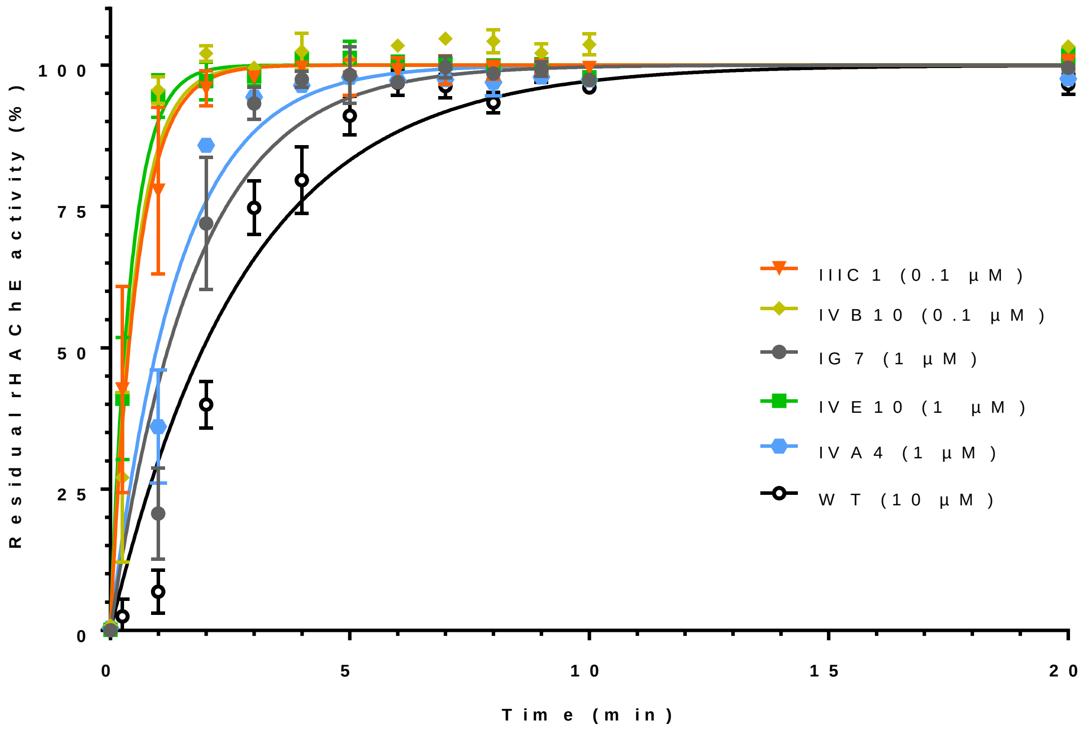

**Figure S5. Protection of rHAChE with *Sso*Pox variants.** rHAChE activity was evaluated after incubation of paraoxon with *Sso*Pox wild type and variants (IIIC1, IVB10, IG7, IVE10, IVA4) for different times. The resulting curves were fitted with *one-phase decay* equation using GraphPad Prism 6 software.

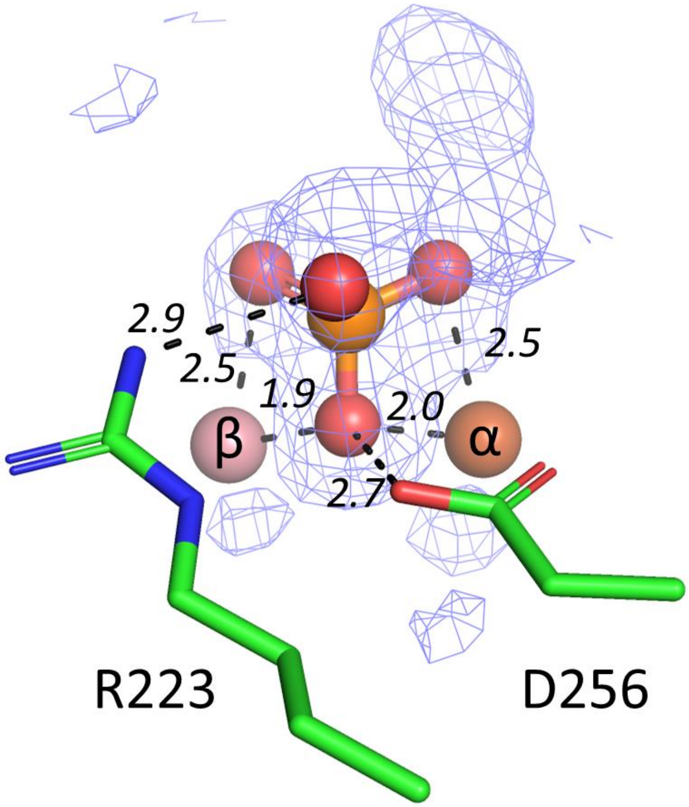

**Figure S6**. **Modeled phosphate anion in the active site of variant IVB10.**

Electronic density is observed in the active sites of several variants, namely IG7, IVA4, IVB10. The active site of variant IVB10 is shown to illustrate this, since the binding is very similar in all variants. The electron density seems to correspond to a tetrahedral molecule that was modelled as a phosphate anion. The omit Fourier difference map $F_{obs}$-$F_{calc}$ is shown as a blue mesh contoured at 2.7σ (carve=3.8) and may indicate the binding of a larger, unidentified molecule. Distances are given in Ångstrom. Metal cations are shown as spheres.

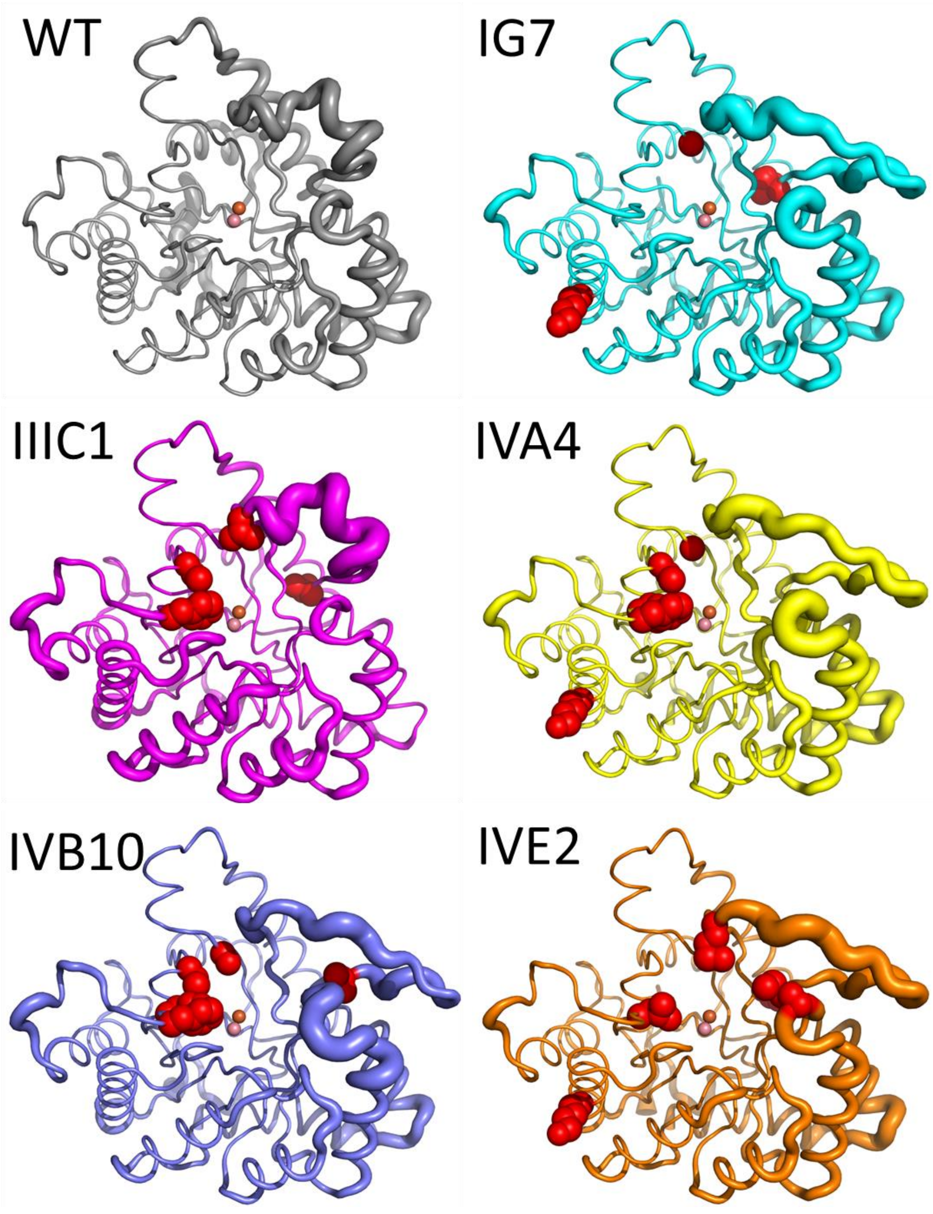

**Figure S7. Structural location of mutations.**

Localization of mutations harbored by *Sso*Pox variants and B-factor putty display for IG7, IIIC1, IVA4, IVB10 and IVE2. Mutations are highlighted in red spheres on each structure. *Sso*Pox wild-type (PDB ID 2VC5) shown for reference.

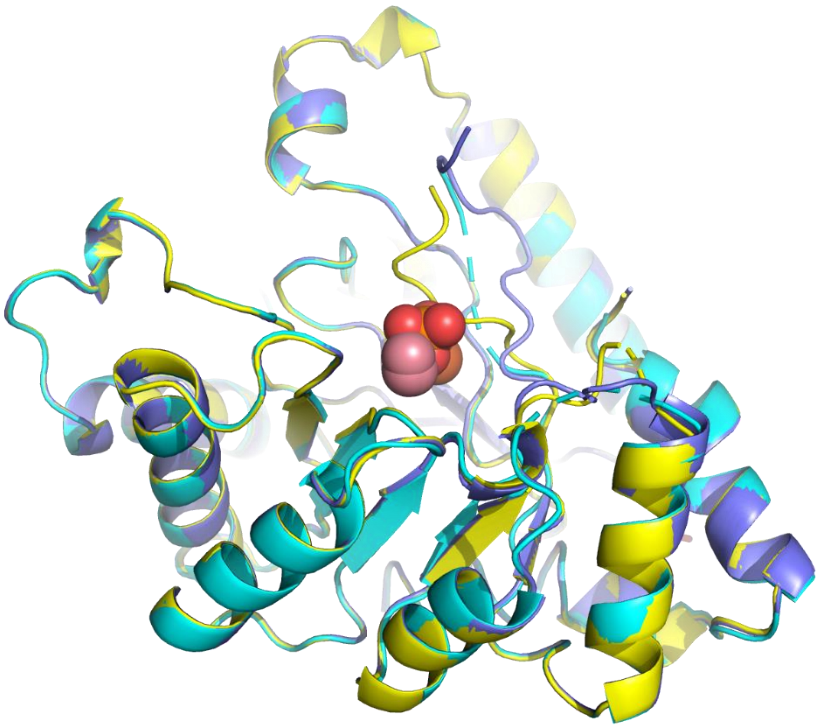

**Figure S8. Superposition of alternative variants' structures.**

The alternative structures of IG7 (cyan), IVB10 (dark blue) and IVA4 (yellow) are superimposed. Modelling for the active site loops 7 and 8 is partial. Metal cations and bound phosphate anion are shown as spheres.

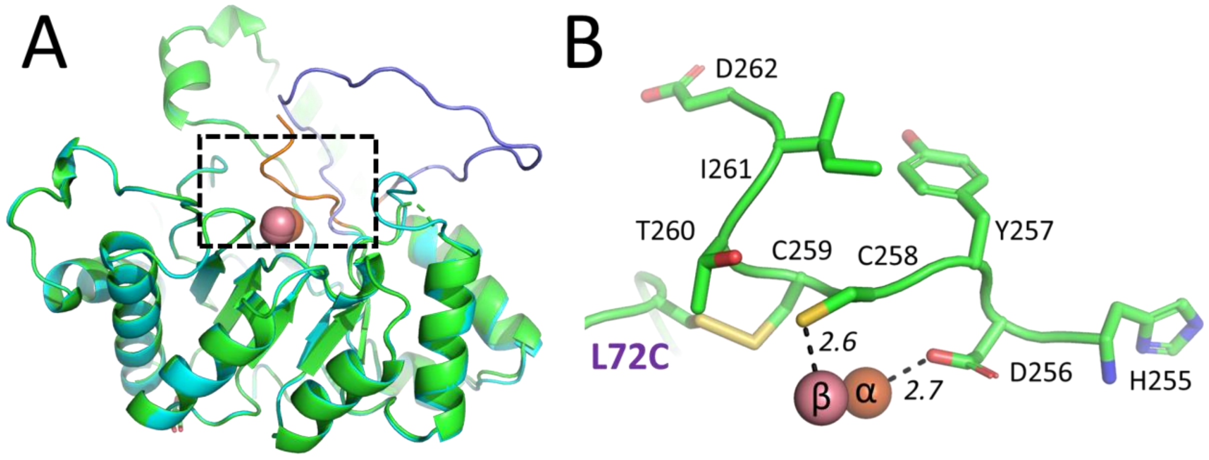

**Figure S9**. **Alternative loop 8 conformation in the second structure of variant IVA4.**

(**A**) The structures obtained for the variant IVA4 are overlaid: the structure with a discrete loop 8 conformation is shown in cyan and blue (loop 8), and the alternative structure is shown in cyan and orange (loop 8). Dashed inset is an approximate location for (**B**) Zoomed-in view on the loop 8 conformation in the IVA4 alternative structure. A disulfide bridge is present between C259 and the mutated residue L72C. C258 is bound to the bimetallic active site, resulting in an inactive enzyme conformation. Distances are given in Ångstrom. Metal cations are shown as spheres.

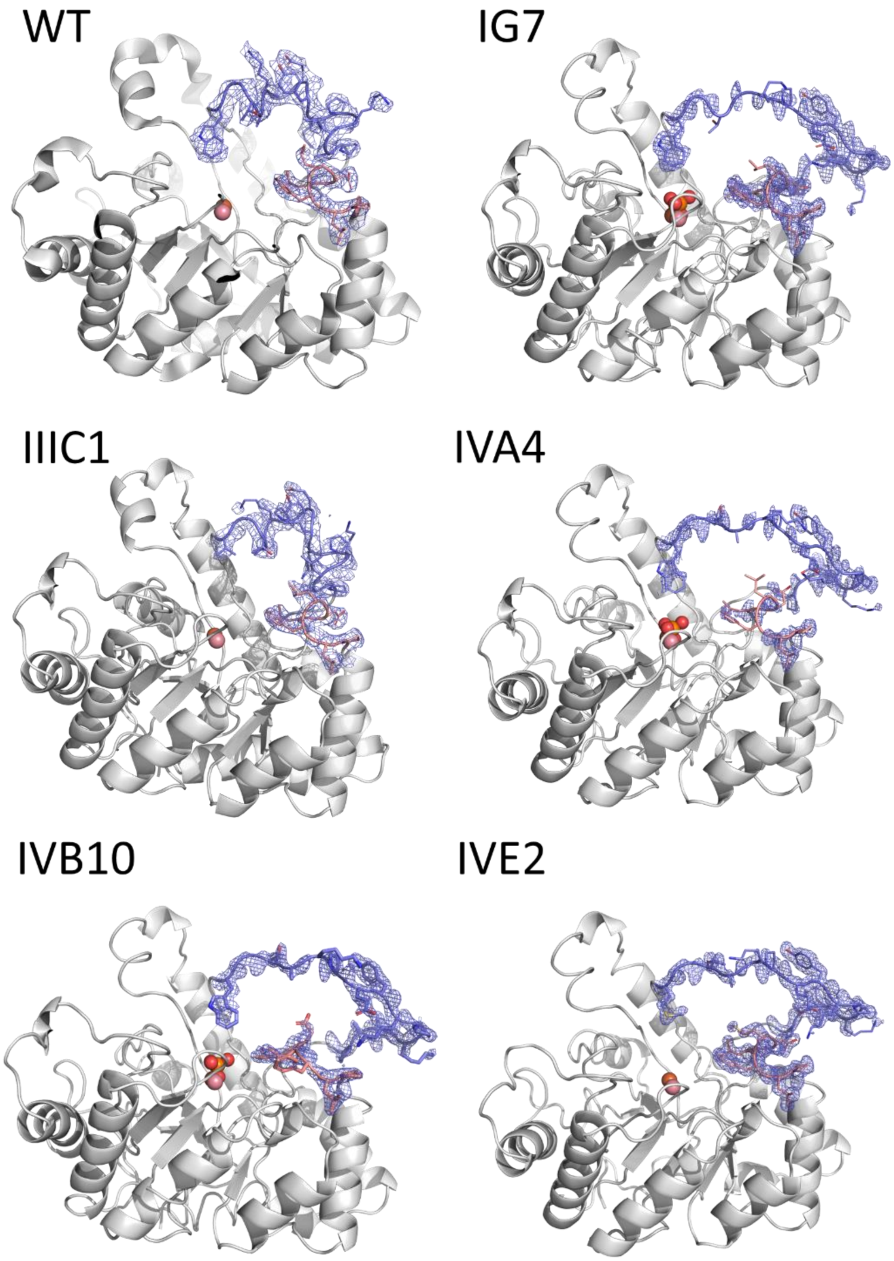

**Figure S10. Electron density maps for loops 7 and 8.**

The $2F_{obs}$-$F_{calc}$ electronic density maps for loop 7 and 8 are shown as blue mesh and contoured at 3σ. Metal cations are shown as spheres.

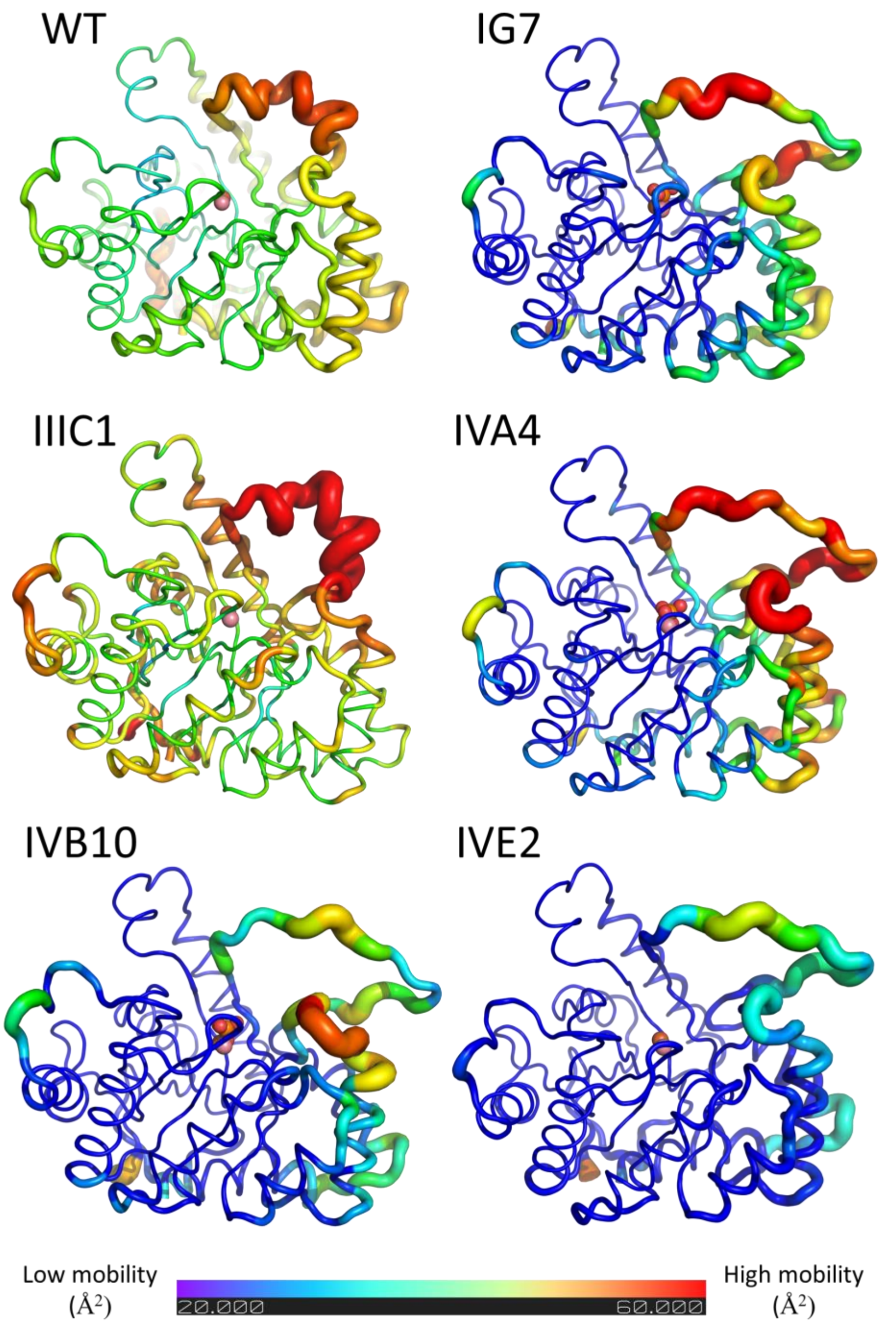

**Figure S11. Representation of thermal motion B-factors.**

Cartoon putty representation of the thermal motion temperature B-factor on the structures of the crystallized *Sso*Pox variants. B-factors are shown on an identical rainbow color scale with dark blue showing the lowest mobility regions (thin) and red the highest mobility regions (thick). Metal cations are shown as spheres.

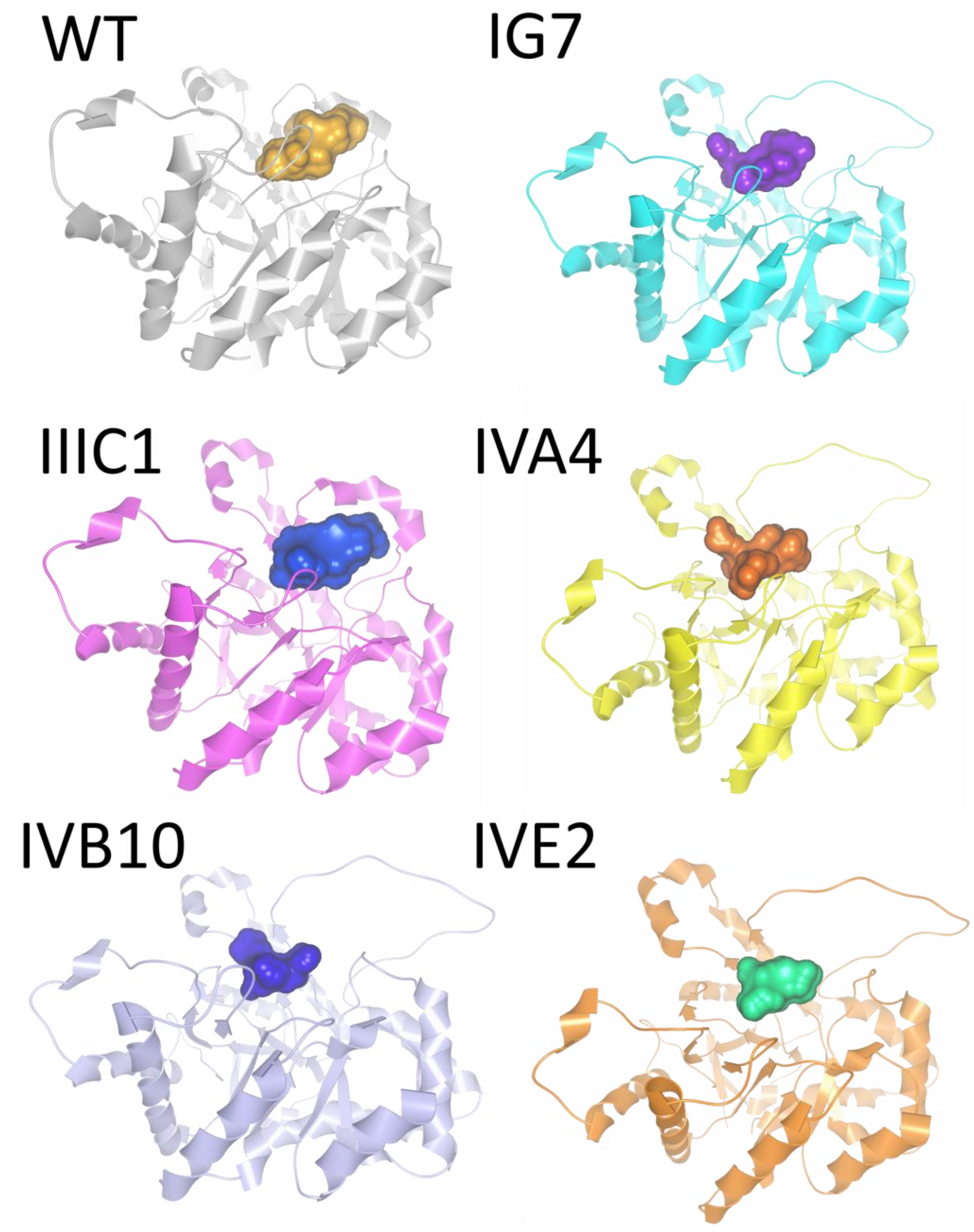

**Figure S12. Active site cavities analysis.**

Active site cavity volumes of the WT enzyme and the crystallized variants using CAVER Analyst 2.0 beta.

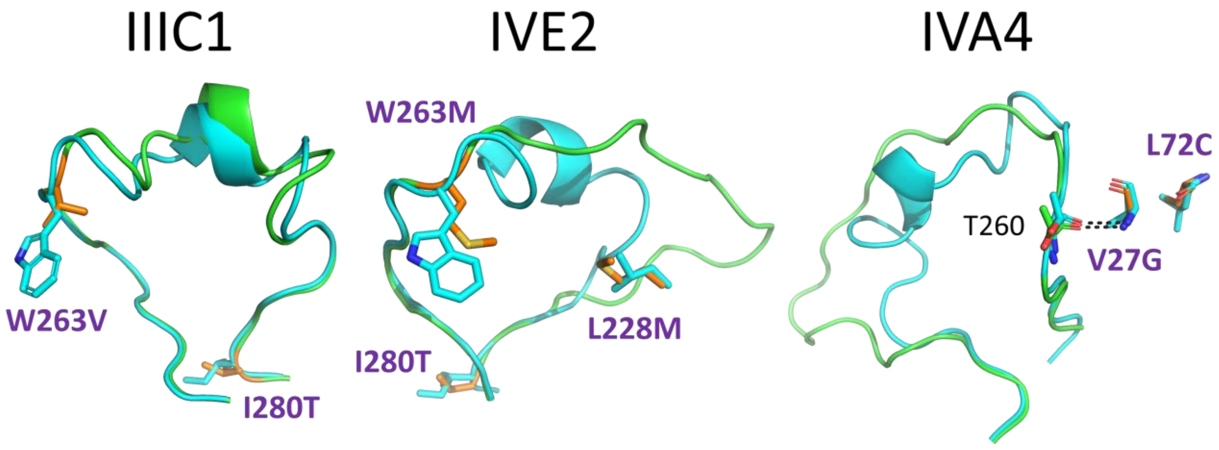

**Figure S13**. **Variant mutations in the vicinity of loop 8.**

Mutated residues are highlighted in orange in the vicinity of the active site loop 8. Structures of the variants (green sticks) are overlaid with the structure of *Sso*Pox-wt (cyan sticks; PDB ID 2VC5). mutation shown in orange.

**Table S1.** Desired mutations and corresponding codons used to synthesize the variants library.

| Residue position | Residue | Codon | Potential mutation | Potential codon | Codon used in oligonucleotides |
|---|---|---|---|---|---|
| 27 | V | GTT | V/A/G | GTT/GCA/GGC | GBT |
| 72 | L | CTG | L/C/I | CTG/TGT/ATT | TGT/MTT |
| 97 | Y | TAT | all | all | NNS |
| 99 | Y | TAT | Y/A/D/E/F | TAT/GCA/GAT/GAG/TTT | KAT/GMA/TTT |
| 154 | R | CGT | R/K | CGT/AAA | ARG |
| 177 | T | ACC | T/D | ACC/GAC | ACC/GAC |
| 223 | R | CGT | all | all | NNS |
| 226 | L | CTG | L/Q/F | CTG/CAG/TTT | TTT/CWG |
| 228 | L | CTG | L/M | CTG/ATG | MTG |
| 258 | C | TGC | C/G/L | TGC/GGC/CTG | KGC/CTG |
| 263 | W | TGG | all | all | NNS |
| 278 | W | TGG | W/L | TGG/CTG | TKG |
| 280 | I | ATT | I/T | ATT/ACT | AYT |

**Table S1. Catalytic parameters of *Sso*Pox variants assayed with organophosphorous compound analogues.** ND, not detected.

| Substrate | *Sso*Pox | $k_{cat}/K_M$ ($M^{-1}.s^{-1}$) | Ratio/*wt* |
|---|---|---|---|
| **Sarin analogue** | *wt* | $3{,}00 \pm 0{,}08 \times 10^2$ | 1 |
| | IH1 | $2{,}16 \pm 0{,}02 \times 10^3$ | 7,20 |
| | IG7 | $2{,}36 \pm 0{,}02 \times 10^2$ | 0,79 |
| | IIIC1 | $5{,}34 \pm 0{,}13 \times 10^3$ | 17,80 |
| | IIIB8 | $9{,}88 \pm 0{,}03 \times 10^1$ | 0,33 |
| | IIIE11 | $3{,}02 \pm 0{,}13 \times 10^4$ | 100,67 |
| | IVE2 | $3{,}90 \pm 0{,}02 \times 10^4$ | 130,00 |
| | IVG2 | $1{,}11 \pm 0{,}002 \times 10^2$ | 0,37 |
| | IVA4 | $4{,}56 \pm 0{,}02 \times 10^2$ | 1,52 |
| | IVE5 | $4{,}79 \pm 0{,}10 \times 10^3$ | 15,97 |
| | IVB10 | $7{,}70 \pm 0{,}12 \times 10^1$ | 0,26 |
| | IVE10 | $9{,}94 \pm 0{,}02 \times 10^2$ | 3,31 |
| | VA7 | $5{,}74 \pm 0{,}01 \times 10^1$ | 0,19 |
| | αsD6 | $9{,}07 \pm 0{,}03 \times 10^3$ | 30,23 |
| | W263I | $2{,}33 \pm 0{,}07 \times 10^4$ | 77,67 |
| **Cyclosarin analogue** | *wt* | $3{,}93 \pm 0{,}20 \times 10^2$ | 1 |
| | IH1 | $1{,}23 \pm 0{,}02 \times 10^3$ | 3,13 |
| | IG7 | $1{,}98 \pm 0{,}08 \times 10^3$ | 5,04 |
| | IIIC1 | $3{,}62 \pm 0{,}07 \times 10^3$ | 9,21 |
| | IIIB8 | $1{,}91 \pm 0{,}01 \times 10^2$ | 0,49 |
| | IIIE11 | $4{,}80 \pm 0{,}08 \times 10^3$ | 12,21 |
| | IVE2 | $4{,}38 \pm 0{,}10 \times 10^5$ | 1114,50 |
| | IVG2 | $8{,}36 \pm 0{,}11 \times 10^2$ | 2,13 |
| | IVA4 | $3{,}84 \pm 0{,}03 \times 10^2$ | 0,98 |
| | IVE5 | $2{,}84 \pm 0{,}10 \times 10^4$ | 72,26 |
| | IVB10 | $1{,}61 \pm 0{,}03 \times 10^2$ | 0,41 |
| | IVE10 | $8{,}19 \pm 0{,}12 \times 10^3$ | 20,84 |
| | VA7 | $1{,}28 \pm 0{,}003 \times 10^2$ | 0,33 |
| | αsD6 | $4{,}93 \pm 0{,}01 \times 10^4$ | 125,45 |
| | W263I | $1{,}89 \pm 0{,}05 \times 10^4$ | 48,09 |
| **Soman analogue** | *wt* | $7{,}63 \pm 0{,}52$ | 1 |
| | IH1 | $7{,}24 \pm 0{,}05 \times 10^1$ | 9,49 |
| | IG7 | ND | ND |
| | IIIC1 | $1{,}37 \pm 0{,}01 \times 10^2$ | 17,96 |
| | IIIB8 | ND | ND |
| | IIIE11 | $1{,}07 \pm 0{,}005 \times 10^2$ | 14,02 |
| | IVE2 | $1{,}11 \pm 0{,}04 \times 10^3$ | 145,48 |
| | IVG2 | ND | ND |
| | IVA4 | $3{,}91 \pm 0{,}02 \times 10^1$ | 5,12 |
| | IVE5 | $8{,}21 \pm 0{,}69 \times 10^3$ | 1076,02 |
| | IVB10 | ND | ND |
| | IVE10 | $2{,}37 \pm 0{,}18 \times 10^3$ | 310,62 |
| | VA7 | ND | ND |
| | αsD6 | $2{,}87 \pm 0{,}02 \times 10^2$ | 37,61 |
| | W263I | $2{,}15 \pm 0{,}02 \times 10^2$ | 28,18 |
| **Tabun analogue** | *wt* | ND | ND |
| | IH1 | ND | ND |
| | IG7 | ND | ND |
| | IIIC1 | $5{,}28 \pm 0{,}11 \times 10^2$ | ND |
| | IIIB8 | ND | ND |
| | IIIE11 | $5{,}09 \pm 0{,}04$ | ND |
| | IVE2 | $7{,}62 \pm 0{,}04$ | ND |
| | IVG2 | ND | ND |
| | IVA4 | $5{,}37 \pm 0{,}05$ | ND |

| | | | |
|---|---|---|---|
| | IVE5 | ND | ND |
| | IVB10 | ND | ND |
| | IVE10 | ND | ND |
| | VA7 | ND | ND |
| | αsD6 | $1,44 \pm 0,01 \times 10^1$ | ND |
| | W263I | $9,22 \pm 0,06 \times 10^1$ | ND |
| **VX analogue** | *wt* | $2,74 \pm 0,02 \times 10^2$ | 1 |
| | IH1 | $4,43 \pm 0,06 \times 10^3$ | 16,17 |
| | IG7 | $3,02 \pm 0,03 \times 10^2$ | 1,10 |
| | IIIC1 | $3,84 \pm 0,21 \times 10^4$ | 140,15 |
| | IIIB8 | $1,45 \pm 0,01 \times 10^2$ | 0,53 |
| | IIIE11 | $9,67 \pm 0,17 \times 10^3$ | 35,29 |
| | IVE2 | $2,15 \pm 0,09 \times 10^5$ | 784,67 |
| | IVG2 | $2,73 \pm 0,02 \times 10^2$ | 1,00 |
| | IVA4 | $2,96 \pm 0,02 \times 10^2$ | 1,08 |
| | IVE5 | $6,80 \pm 0,10 \times 10^3$ | 24,82 |
| | IVB10 | $6,68 \pm 0,06 \times 10^2$ | 2,44 |
| | IVE10 | $7,28 \pm 1,22 \times 10^3$ | 26,57 |
| | VA7 | $8,41 \pm 0,02 \times 10^1$ | 0,31 |
| | αsD6 | $3,91 \pm 0,08 \times 10^3$ | 14,27 |
| | W263I | $8,15 \pm 0,33 \times 10^4$ | 297,45 |

**Table S2.** Data collection and structure refinement statistics.

| | IG7 | IIIC1 | IVA4 | IVB10 | IVE2 |
|---|---|---|---|---|---|
| PDB ID | 8SF2 | 8SFA | 8SFB | 8SFD | 8SFK |
| Beamline | APS 23ID-B | | | | |
| Detector | Dectris Eiger-16M | | | | |
| Wavelength (Å) | 0.96802 | 1.033202 | 1.03321 | 1.03321 | 0.96802 |
| Resolution (Å) | 1.5 | 2.32 | 1.4 | 1.5 | 1.4 |
| Space group | $C222_1$ | $P2_12_12_1$ | $C222_1$ | $C222_1$ | $C222_1$ |
| *Unit cell* dimensions (Å) | a= 65.4 b=74.7 c=137.5 | a= 90.2 b=104.3 c=153.8 | a= 64.5 b=74.9 c=137.6 | a= 65.3 b=74.7 c=138.7 | a= 66.1 b=74.7 c=137.4 |
| angles (°) | α = β = γ = 90° | | | | |
| No. observed reflections (last bin) | 303371 (54456) | 507054 (62600) | 492631 (71617) | 450128 (79833) | 477153 (85133) |
| No. unique reflections (last bin) | 53878 (9360) | 62986 (7364) | 65450 (11902) | 54520 (9490) | 67061 (12397) |
| Completeness (%) (last bin) | 99.4 (99.3) | 99.2 (98.9) | 99.6 (97.7) | 100 (100) | 100 (99.9) |
| $R_{meas}$ (%) (last bin) | 3.6 (33.0) | 6.8 (53.6) | 3.6 (59.5) | 5.5 (59.9) | 6.3 (63.3) |
| $CC_{1/2}$ | 100.0 (97.7) | 99.9 (97.8) | 100.0 (92.2) | 99.9 (94.5) | 100.0 (91.7) |
| I/σ(I) (last bin) | 26.70 (2.87) | 19.78 (4.54) | 27.53 (2.94) | 21.5 (3.95) | 19.91 (3.26) |
| Last resolution shell | 1.6 – 1.5 | 2.42 - 2.32 | 1.5 – 1.4 | 1.6 -1.5 | 1.5 – 1.4 |
| Redundancy (last bin) | 5.63 (5.82) | 8.05 (8.5) | 7.5 (6.01) | 8.25 (8.41) | 4.34 (5.61) |
| Resolution | 49.2 - 1.5 | 68.23 - 2.32 | 68.78 - 1.40 | 49.14 - 1.50 | 49.47 - 1.40 |
| $R_{free}/R_{work}$ | 18.98/15.87 | 24.19/19.01 | 22.26/19.99 | 17.27/14.02 | 16.93/13.54 |
| rmsd bond length (Å) | 0.005 | 0.008 | 0.005 | 0.005 | 0.005 |
| rmsd bond angle (°) | 1.012 | 0.894 | 1.048 | 0.846 | 0.869 |
| Ramachandran outliers (%) | 0.00 | 0.16 | 0.65 | 0.00 | 0.00 |
| Ramachandran favored (%) | 96.44 | 96.28 | 96.44 | 96.12 | 97.41 |

**Table S4.** Data collection and structure refinement statistics for alternate, disordered conformations.

| | IG7 | IVA4 | IVB10 |
|---|---|---|---|
| PDB ID | 8SF9 | 8SFC | 8SFM |
| Beamline | APS 23ID-B | | |
| Detector | Dectris Eiger-16M | | |
| Wavelength (Å) | 0.774901 | 0.774901 | 0.774901 |
| Resolution (Å) | 1.8 | 1.4 | 1.5 |
| Space group | $C222_1$ | $C222_1$ | $C222_1$ |
| Unit cell dimensions (Å) | a = 64.52<br>b = 74.96<br>c = 137.30 | a = 64.09<br>b = 74.95<br>c = 137.11 | a = 64.11<br>b = 74.61<br>c = 137.69 |
| angles (°) | α = β = γ = 90° | α = β = γ = 90° | α = β = γ = 90° |
| No. observed reflections (last bin) | 202464 (31855) | 418892 (78088) | 378951 (67696) |
| No. unique reflections (last bin) | 58954 (8881) | 124395 (23305) | 101746 (17921) |
| Completeness (%) (last bin) | 98.8 (99.2) | 98.9 (99.0) | 99.6 (99.6) |
| $R_{meas}$ (%) (last bin) | 5.8 (77.0) | 4.4 (68.2) | 6.9 (46.1) |
| $CC_{1/2}$ | 99.8 (83.2) | 99.9 (83.0) | 99.7 (87.4) |
| I/σ(I) (last bin) | 11.34 (2.09) | 15.22 (2.32) | 11.50 (3.17) |
| Last resolution shell | 1.9-1.8 | 1.5-1.4 | 1.6-1.5 |
| Redundancy (last bin) | 3.43 (3.58) | 3.36 (3.35) | 3.72 (3.77) |
| Resolution | 46.07 - 1.80 | 37.47 - 1.40 | 34.42 - 1.50 |
| $R_{free}/R_{work}$ | 23.37/20.04 | 22.59/20.48 | 23.03/20.23 |
| rmsd bond length (Å) | 0.004 | 0.007 | 0.007 |

| | | | |
|---|---|---|---|
| rmsd bond angle (°) | 0.65 | 0.87 | 0.92 |
| Ramachandran outliers (%) | 0.00 | 1.06 | 0.00 |
| Ramachandran favored (%) | 93.80 | 95.76 | 96.17 |

**Table S5.** Mutation list of the selected *Sso*Pox variants and their corresponding melting temperature values $T_m$ (°C). Data for wt, asD6 and W263 are taken from previous work[1,2,3].

| Variant | Mutations | $T_m$ (°C) |
|---|---|---|
| **wt**[1] | - | 106 |
| **IH1** | Y97Q/Y99F | 86,2 ± 0,3 |
| **IG7** | V27G/R154K/I280T | 79,1 ± 1,7 |
| **IIIC1** | L72C/Y97F/Y99F/W263V/I280T | 96,3 ± 0,5 |
| **IIIB8** | L72I/Y97W/Y99F/W278L/I280T | 79,8 ± 0,1 |
| **IIIE11** | Y97S/Y99F | 84,1 ± 2,7 |
| **IVE2** | Y97L/R154K/L228M/W263M/I280T | 89,7 ± 0,5 |
| **IVG2** | V27A/L72C/Y97R/Y99A/I280T | 79,2 ± 0,1 |
| **IVA4** | V27G/L72C/Y97I/Y99F/R154K | 86,7 ± 0,2 |
| **IVE5** | Y97F/Y99F/R154K/C258L/W263D/I280T | 82,3 ± 0,1 |
| **IVB10** | V27A/L72C/Y97F/Y99F/W278L | 85,0 ± 0,2 |
| **IVE10** | V27A/Y97F/Y99F/R154K/T177D/W263F | 81,5 ± 0,1 |
| **VA7** | V27A/Y99D | 82,3 ± 0,3 |
| **αsD6**[2] | V27A/Y97W/L228M/W263M | 82,5 ± 1,8 |
| **W263I**[3] | W263I | 87,8 ± 1,2 |

### References.

(1) Merone, L.; Mandrich, L.; Rossi, M.; Manco, G. A Thermostable Phosphotriesterase from the Archaeon Sulfolobus Solfataricus: Cloning, Overexpression and Properties. *Extremophiles* **2005**, *9* (4), 297–305. https://doi.org/10.1007/s00792-005-0445-4.
(2) Jacquet, P.; Hiblot, J.; Daudé, D.; Bergonzi, C.; Gotthard, G.; Armstrong, N.; Chabrière, E.; Elias, M. Rational Engineering of a Native Hyperthermostable Lactonase into a Broad Spectrum Phosphotriesterase. *Scientific Reports* **2017**, *7* (1). https://doi.org/10.1038/s41598-017-16841-0.
(3) Hiblot, J.; Gotthard, G.; Elias, M.; Chabriere, E. Differential Active Site Loop Conformations Mediate Promiscuous Activities in the Lactonase SsoPox. *PLoS ONE* **2013**, *8* (9), e75272. https://doi.org/10.1371/journal.pone.0075272.
